# UTag, a cysteine-free thermostable tagging system for tracking single mRNA translation live

**DOI:** 10.64898/2026.05.06.723082

**Authors:** Luis U Aguilera, Szuhsuan Chen, Rhiannon M. Sears, Jake Yarbro, Jacob DeRoo, Hunter A. Ogg, Brian J. Geiss, Timothy J. Stasevich, Christopher D. Snow, Ning Zhao

**Affiliations:** Department of Biochemistry and Molecular Genetics, University of Colorado-Anschutz Medical Campus, Aurora, CO, 80045, USA; School of Biomedical and Chemical Engineering, Colorado State University, Fort Collins, CO, 80523, USA; Department of Microbiology, Immunology, and Pathology, Colorado State University, Fort Collins, CO, 80523, USA; Department of Biochemistry and Molecular Biology, Colorado State University, Fort Collins, CO, 80523, USA

## Abstract

Spatiotemporal regulation of mRNA translation is central to gene expression. Over the past decade, translation has become directly observable in live cells at single-mRNA resolution by tagging nascent chains with tandem arrays of short epitope tags recognized by genetically encodable fluorescent intracellular antibodies (intrabodies). While this technology has revolutionized our understanding of translation regulation, the current toolbox of tagging systems remains limited. Here, we developed a novel and tight-binding intrabody against a short (11-amino acid) HIV protease epitope (named UTag). To ensure robust intracellular folding of the anti-UTag intrabody, we further engineered a cysteine-free variant that folds and functions independently of disulfide-bond formation, as validated by X-ray crystallography. The cysteine-free anti-UTag intrabody retains high binding affinity comparable to the parental intrabody while exhibiting significantly improved thermostability (∼80 °C). Importantly, the cysteine-free UTag system enables real-time tracking of single-mRNA translation in live cells with performance on par with the parental UTag system as well as the established SunTag and ALFA-tag. Collectively, these results demonstrate that the newly developed UTag system expands the toolbox for live-cell translation tracking and provides complementary tools for multiplexed applications.

## Introduction

mRNA translation within the central dogma is spatiotemporally regulated to maintain protein homeostasis and cellular function. Translation regulation has been intensively studied by conventional biochemical approaches, such as western blotting, luciferase assays, and fluorescent protein reporters. However, these approaches measure total protein output and cannot distinguish newly synthesized proteins from pre-existing pools, thereby obscuring the regulation of the true translation process. Moreover, these bulk assays inherently overlook the spatially localized and stochastic kinetics of individual translation events.^1^ Over the past decade, translation has become directly observable in live cells through real-time tracking of nascent chains at single-mRNA level.^2–6^ This technology has transformed our understanding of translation regulation, revealing layers of complexity, including kinetic heterogeneity among mRNAs^7^, subcellularly localized translation variability^4,8,9^, stochastic translational bursting^10^, ribosome cooperativity^11^ and collisions^12^, translation under cellular stress^13,14^, and the kinetics of noncanonical translation, such as frameshifting^15^ as well as internal ribosome entry site (IRES)-^16^ or repeat-associated non-AUG (RAN)-initiated translation^17^.

To track single-mRNA translation in live cells, a tandem array (10-56x) of short epitope tags is fused to the N- or C-terminus of a protein of interest (POI).^2–6,18–20^ These tags enable direct and continuous visualization of nascent chains through binding by cognate genetically encodable fluorescent intracellular antibodies (intrabodies), including single-chain antibody fragments (scFvs) and nanobodies (nbs). Since translation was first directly visualized using this strategy, significant effort has been devoted to expanding the toolbox for translation tracking^2,7,18,21–24^. Several key factors guide the development of suitable tagging systems: (1) epitope tags should be small to minimize the overall size of tandem tags and reduce perturbation to the POI; (2) cognate intrabodies must fold properly in live cells to enable high-specificity and high-affinity binding to their epitope tags; (3) linkers between tandem tags must be optimized to achieve maximal intrabody binding stoichiometry; and (4) for multiplexed imaging, newly developed tags should be comparable in performance and orthogonal to existing time-tested tagging systems.

Current tagging systems, including smHA^22,25^, smFLAG^23,25^, SunTag^24^, MoonTag^7^, and ALFA-tag^18^, have been developed following these design strategies. The smHA and smFLAG tags consist of 10 linear, unstructured HA or FLAG epitopes embedded within loops of mRuby or sfGFP, enabling recruitment of up to 10 fluorescent intrabodies.^22,23,25^ In contrast, SunTag, MoonTag, and ALFA-tag form an α-helical secondary structure and can be conveniently assembled into large tandem tags using flexible linkers.^5,7,18^ The tandem arrays of 24x, 32x, or even 56x tags have been implemented to enhance signal-to-noise at translation sites.^3,5,18^ These tagging systems span a broad range of effective binding affinities, determined by both the intrinsic affinity of individual epitope tags and intrabodies (nM∼pM) and the avidity effect conferred by repeated tag arrays. A key consideration across these systems is the balance between binding affinity and perturbation of the tagged protein: lower-affinity interactions can limit signal-to-noise (e.g. smHA, smFLAG, and MoonTag), whereas extremely high-affinity binding enhances signal but may, in some contexts, influence protein behavior or localization. This tradeoff highlights the need for tagging strategies with favored high-affinity binding that provide robust signal while preserving native protein localization and function. Thus, there remains a need to develop novel tagging systems that are comparable in performance and orthogonal to established bright tagging systems, such as SunTag and ALFA-tag, thereby enabling robust multiplexed study of translation kinetics.

One major bottleneck in developing such tagging systems is the difficulty intrabodies have when folding in the reducing live cell environment. This difficulty arises from the fundamental biology of antibody biogenesis. Full-length antibodies are synthesized, folded, and assembled in the endoplasmic reticulum (ER), where an oxidizing environment promotes disulfide bond formation that is critical for proper folding and assembly.^26,27^ In contrast, intrabodies, which consist of one (nanobodies) or two (scFvs) variable domains of full-length antibodies, are synthesized and folded in the cytosol where a reducing environment is maintained.^28,29^ As a result, disulfide-bonds in intrabodies cannot be formed robustly, often leading to misfolding and loss of epitope-binding capacity.^21–23^ To overcome this intracellular folding challenge, protein engineering has been employed to improve intrabody folding. However, these approaches typically require extensive screening of variants, limiting throughput and efficiency. More recently, advances in artificial intelligence have significantly accelerated this process.^30^ Complementary to these approaches, we addressed this challenge by grafting the complementarity determining region (CDR) loops of scFvs that misfold in live cells onto stable scFv scaffolds that have been validated for live-cell imaging and are therefore less dependent on disulfide bond formation.^22,23^

In this study, we leveraged the previously identified stable scaffold, 15F11, to mine antibody fragments deposited in the Protein Data Bank (PDB) and ranked candidates based on sequence-homology. Through this approach, we identified a fragment antigen-binding (Fab) targeting HIV protease^31^ that shares ∼85% sequence identity with the 15F11 scaffold. Its binding epitope is an 11-amino acid (aa) peptide (MSLPGRWKPKM) that adopts a stable U-shaped secondary structure and is well suited for tandem repetition using flexible linkers.^31^ We therefore designated this novel epitope as “UTag”, reflecting its distinctive structural feature. By grafting the CDR loops of this Fab onto the 15F11 scaffold, we developed a new anti-UTag intrabody that binds its epitope with high specificity and affinity in live cells. We further engineered this molecule to remove all cysteine residues, yielding a cysteine-free variant that exhibits enhanced thermostability while retaining tight binding to UTag, independent of disulfide bond formation. We demonstrate that the cysteine-free intrabody enables single-mRNA translation tracking in live cells, with fluorescence intensity and translation kinetics comparable to the parental intrabody and established SunTag and ALFA-tag systems. Together, the newly developed UTag system expands the current toolbox for translation tracking and provides a platform for multiplexed tracking in combination with SunTag and ALFA-tag systems.

## Results

### Development and initial screening of anti-UTag intrabodies

scFvs can be conveniently engineered by fusing the variable regions of antibody heavy and light chains with a flexible linker. When properly folded, scFvs preserve the binding affinity and specificity of their parental antibodies and can function as genetically encoded intrabodies for live-cell imaging. The increasing availability of antibody sequences has greatly expanded the scFv pool, accelerating the development of intrabody-based imaging tools. In practice, however, most scFvs fail to fold correctly in the intracellular environment, which substantially limits their application in live cells.

To rescue scFv folding and function in live cells, engineering their scaffold is usually required. Previously, we identified two stable scFv scaffolds, 15F11 (**Fig. 1A**; PDB: 5B3N) and 2E2, that fold properly in live cells. Grafting the CDR loops of anti-HA and anti-FLAG scFvs onto these scaffolds successfully rescued their folding and restored binding to their cognate HA and FLAG tags in live cells.^22,23^ To expand the toolbox of intrabody-based probes for live-cell imaging, we used the stable 15F11 scaffold to mine antibody fragments from the PDB and ranked candidates based on sequence identity to the 15F11 scaffold. This analysis identified a Fab (∼85% sequence identity) that binds a short, U-shaped 11-aa epitope (coined UTag) derived from HIV protease^31^ (**Fig. 1B**; PDB: 2HRP).

**Figure 1.**
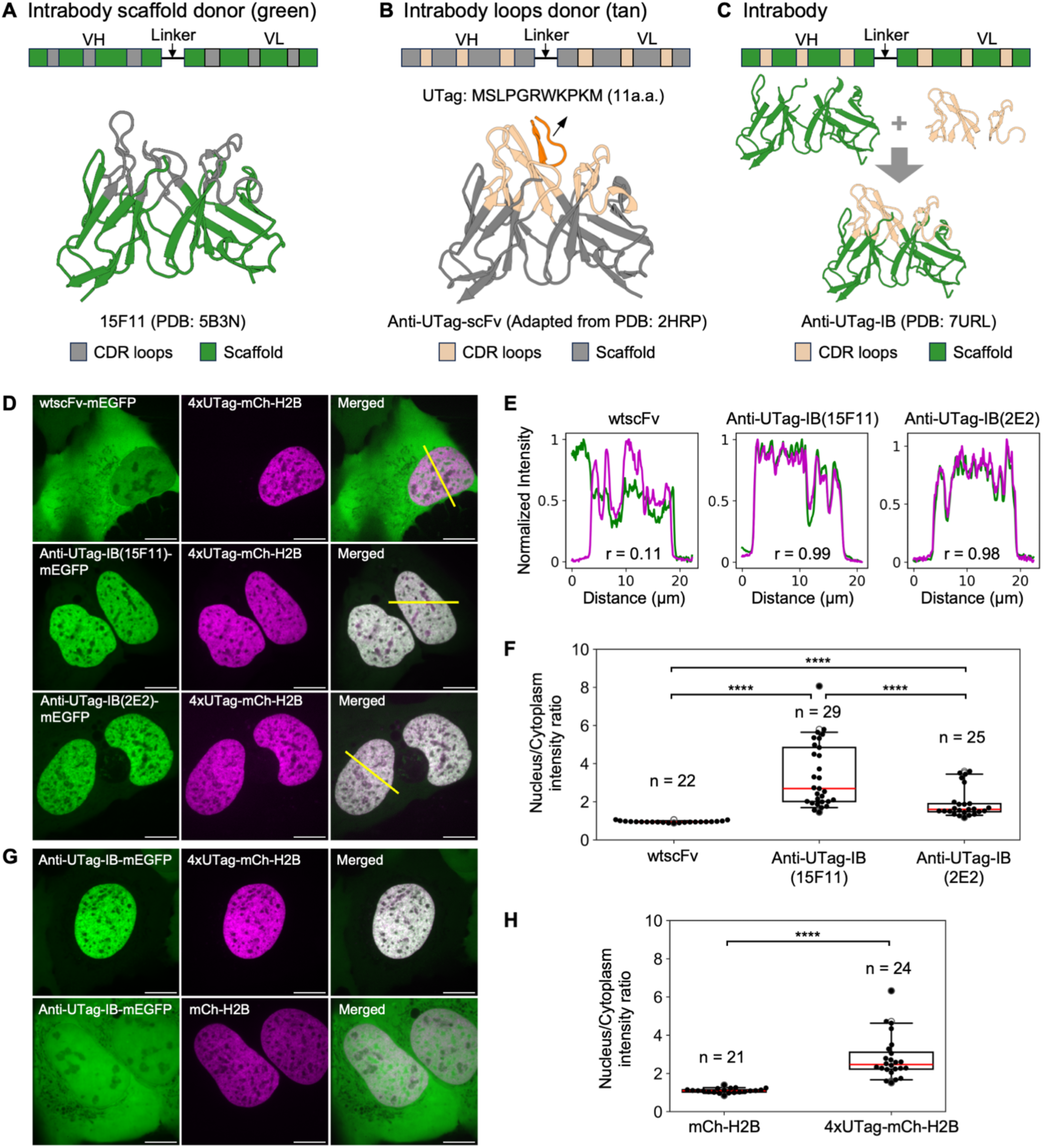
Development of anti-UTag intrabodies and initial screening results. **(A-C)** Schematics illustrating the scaffold (PDB: 5B3N; A, green) and CDR loops (PDB: 2HRP; B, tan) used to generate the anti-UTag intrabody (anti-UTag-IB; PDB: 7URL; C). **(D)** Representative images of cells co-expressing mEGFP-tagged wild-type anti-UTag scFv (wtscFv-mEGFP, top), anti-UTag intrabody generated using the 15F11 scaffold (anti-UTag-IB(15F11)-mEGFP, middle), or the 2E2 scaffold (anti-UTag-IB(2E2)-mEGFP, bottom), together with 4xUTag-mCh-H2B. **(E)** Normalized fluorescence intensity profiles along the yellow lines shown in D for both green and magenta channels. Intensities were normalized by setting the minimum and maximum values to 0 and 1, respectively. Pearson correlation coefficients (r) are indicated. **(F)** Nuclear-to-cytoplasmic mEGFP fluorescence intensity ratios for all cells imaged in D. **(G)** Representative images of cells co-expressing anti-UTag-IB(15F11)-mEGFP with either 4xUTag-mCh-H2B (top) or mCh-H2B (bottom). **(H)** Nuclear-to-cytoplasmic mEGFP fluorescence intensity ratios for all cells imaged in G. Mann-Whitney U test. **** p < 0.0001. Scale bars: 10 µm. In box-and-whisker plots, red lines indicate medians.

We first generated a wild-type anti-UTag scFv by fusing the variable regions of the Fab heavy and light chains with a flexible linker, as described previously^22,23,32^ (**Fig. 1B**). To assess its ability to bind UTag in live cells, we performed a live-cell co-localization assay by co-expressing mEGFP-tagged wild-type scFv (wtscFv-mEGFP) with 4xUTag-tagged mCh-H2B (4xUTag-mCh-H2B) in U-2 OS cells. If the scFv folds correctly, it should retain its ability to bind UTag and co-localize with 4xUTag-mCh-H2B in the cell nucleus. However, the wtscFv-mEGFP displayed a diffuse, predominantly cytosolic distribution without detectable nuclear enrichment (**Fig. 1D**, top) and showed little correlation with the mCh signal (**Fig. 1E**, left; Pearson’s r = 0.11). These results indicate that the wild-type anti-UTag scFv cannot bind UTag in live cells, likely due to misfolding in the reducing intracellular environment. To rescue its intracellular folding, we grafted all six CDR loops of the wild-type anti-UTag scFv (**Fig. 1B**, tan) onto either the 15F11 or 2E2 stable scaffolds (**Fig. 1A**, green), generating two anti-UTag intrabodies (**Fig. 1C**). To evaluate their binding to UTag, each intrabody was fused to mEGFP and tested using the same live-cell co-localization assay. Excitingly, both intrabodies exhibited clear nuclear enrichment when co-expressed with 4xUTag-mCh-H2B (**Fig. 1D**, middle and bottom) and showed strong correlation with the mCh signal (Pearson’s r = 0.99 and 0.98, respectively) (**Fig. 1E**, middle and right). Quantitative analysis of the nucleus-to-cytoplasmic mEGFP fluorescence intensity ratios further confirmed the robust nuclear enrichment of both intrabodies (**Fig. 1F**; p < 0.0001). Because the 15F11-grafted intrabody showed significantly higher nuclear enrichment than the 2E2-grafted one, we selected the 15F11-grafted intrabody for all subsequent characterization, hereafter referred to as anti-UTag intrabody (anti-UTag-IB).

To confirm that the nuclear localization of the intrabody resulted from specific UTag binding rather than nonspecific effects, we performed a control experiment in which the UTag epitope was omitted. In cells co-expressing anti-UTag-IB-mEGFP with 4xUTag-mCh-H2B, the intrabody localized to the nucleus as expected (**Fig. 1G**, top). In contrast, when co-expressed with mCh-H2B lacking the UTag epitope, the intrabody remained diffuse and was evenly distributed between the cytoplasm and nucleus (**Fig. 1G**, bottom). Consistent with this observation, the nuclear-to-cytoplasmic fluorescence intensity ratios of mEGFP was significantly higher when co-expressed with 4xUTag-mCh-H2B (2.82 ± 0.24, mean ± SEM) than with mCh-H2B (1.10 ± 0.03, mean ± SEM) (**Fig. 1H**; p < 0.0001), corresponding to a ∼2.6-fold enrichment. These results confirm that nuclear accumulation of the intrabody depends on specific interaction with UTag in live cells.

### Crystal structure reveals that anti-UTag intrabody retains the UTag-binding pocket with reduced folding dependence on disulfide bonds

To ensure that loop grafting onto the 15F11 scaffold did not distort the UTag-binding pocket, we solved its crystal structure (PDB: 7URL) at 1.49 Å resolution. As shown in **Fig. 2A**, the intrabody structure closely superimposes on the anti-UTag Fab variable regions (PDB: 1MF2; RMSD_Cα_ = 0.8 Å), demonstrating that the loop grafting process preserves the conformation of the CDR loops and, consequently, the integrity of the UTag-binding pocket.

**Figure 2.**
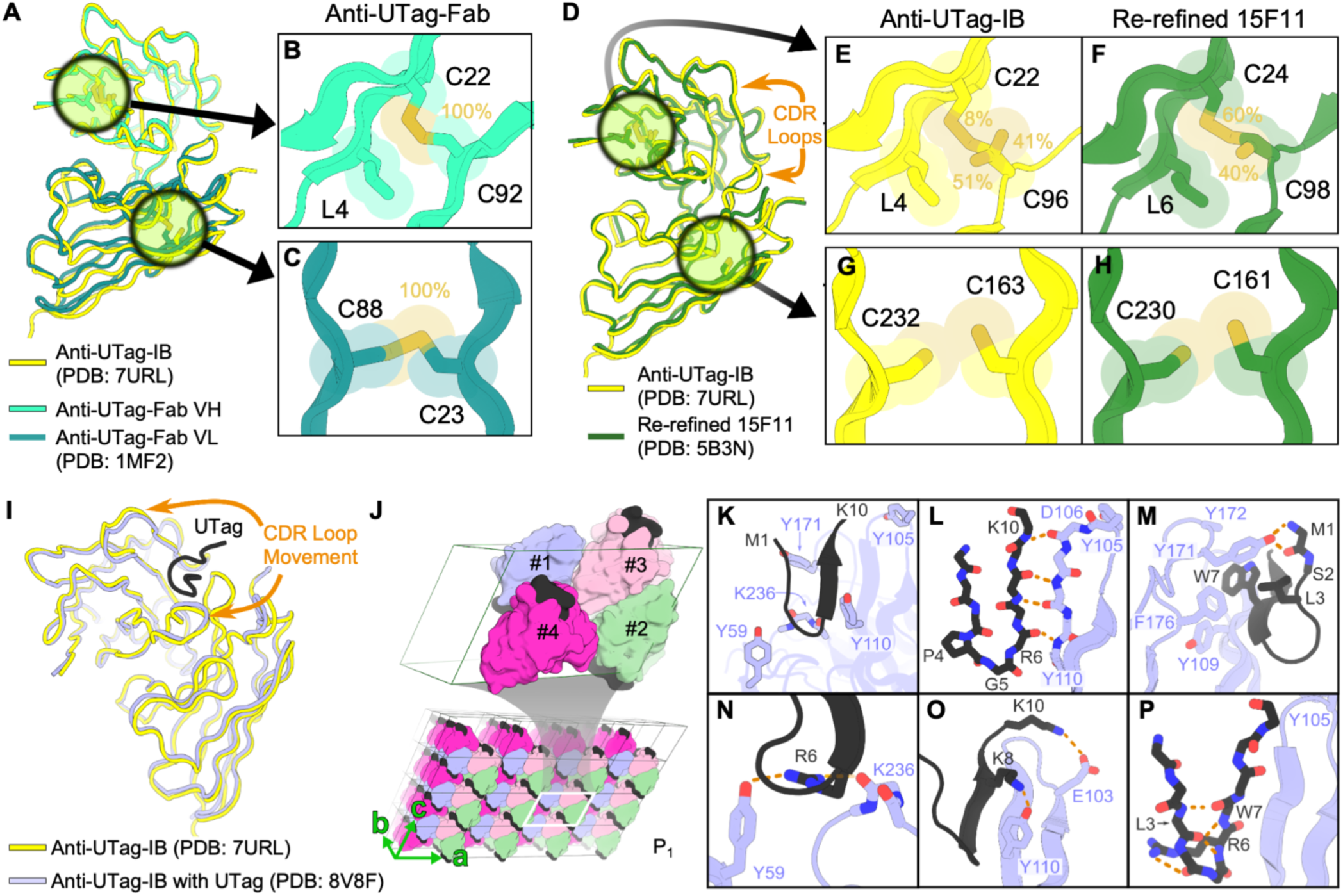
Structural characterization and crystal lattice analysis of anti-UTag-IB. **(A)** Structural superposition of anti-UTag-IB and anti-UTag-Fab, showing the separate heavy chain (top, cyan-green) and light chain (bottom, teal). RMSD_Cα_ = 0.8 Å. **(B-C)** Detailed views of the disulfide bonds between heavy chain C22 and C92, and light chain C23 and C88 in anti-UTag-Fab. Occupancy percentages for side-chain conformations are indicated. **(D)** Structural superposition of anti-UTag-IB and re-refined 15F11 scFv. RMSD_Cα_ = 0.9 Å. **(E-F)** Detailed view of the C22-C96 disulfide bond (C24-C98 in re-refined 15F11). Occupancy percentages for alternate side-chain conformations in the intrabody (E) and the re-refined 15F11 (F) are indicated. **(G-H)** Detailed view of the C163-C232 disulfide bond (C161-C230 in re-refined 15F11), confirming no disulfide-bond formation in both intrabody (G) and re-refined 15F11 (H). **(I)** Structure alignment of anti-UTag-IB with the UTag-bound complex. RMSD_Cα_ = 0.7 Å. **(J)** Crystal packing arrangement of the intrabody-UTag complex (PDB: 8V8F). The P_1_ unit cell contains four independent copies in the asymmetric unit (blue, green, pink, and magenta). A 3×3x3 supercell expansion is shown with crystallographic axes a, b, and c indicated. **(K–P)** Representative views of interactions from Copy 1 in J. (**K**) Overview of the UTag peptide (black) and key intrabody residues (blue) involved in inter-chain interactions. **(L–O)** Inter-chain interactions between UTag (black) and intrabody CDR loops (blue). **(P)** Intra-chain sidechain-to-backbone and backbone-to-backbone interactions that stabilize the UTag β-hairpin. Hydrogen bonds and salt bridges are indicated by dashed lines, with participating residues labeled.

However, the two disulfide bonds in the Fab variable regions: C22-C92 in the heavy chain and C23-C88 pair in the light chain (**Figs. 2B-2C**; PDB: 1MF2), are not likely to form in the intrabody. At high resolution (1.49 Å), we observed no evidence of disulfide bond formation for C163-C232 (equivalent to C23-C88 in the Fab light chain), while C22-C96 (equivalent to C22-C92 in the Fab heavy chain) shows only ∼8% occupancy for rotamers consistent with disulfide bonding (**Figs. 2E** and **2G**; PDB: 7URL). Reduced folding dependence on disulfide bonds may be inherited from the scaffold donor 15F11. To corroborate this hypothesis, we re-refined the existing 15F11 crystal structure (PDB: 5B3N). Re-refined 15F11 superimposed on the intrabody with a RMSD_Cα_ = 0.9 Å (**Fig. 2D**), confirming scaffold preservation. The improved electron density map was consistent with low disulfide yields of ∼60% and 0%, respectively (**Figs. 2F, 2H,** and **S1**). Furthermore, the re-refined model exhibited improvement in model validation statistics reported from wwPDB validation tool (**Table S1**), including a reduction in R_free_ (from 0.253 to 0.234) and a drop in the Clash score from 3 to 1. Notably, sidechain outliers were eliminated entirely (reducing from 2.1% to 0), ensuring a higher-quality model to support our observations. Together, our re-refined model validates that the 15F11 folds with significantly reduced dependence on disulfide-bonds, confirming that high intrabody performance in the absence of disulfide bonds is inherited from the 15F11 scaffold.

To further characterize the interaction between the intrabody and UTag, we determined the co-crystal structure of the complex (PDB: 8V8F) at 2.27 Å resolution. Despite the use of similar growth conditions, the complex crystallized in the P_1_ space group rather than the P_212121_ space group observed for the apo-intrabody (PDB: 7URL) (**Fig. S2**; **Tables S3** and **S5**). Comparison of the two structures revealed a subtle CDR loop movement (RMSD_Cα_ = 0.7 Å) upon UTag binding, as shown in **Fig. 2I**.

Structural analysis of the four copies in the unit cells (**Fig. 2J**) for the anti-UTag-IB/UTag complex (PDB: 8V8F) reveals a robust network of inter-chain interactions that stabilize the UTag within the anti-UTag-IB binding groove (**Fig. 2K**), that are consistent with the parental CDR-donor structure (PDB: 2HRP). The UTag structure is centered around a type II β-turn formed by residues P4 (Φ = −55.3° ± 4.8°, Ψ = 130.4° ± 0.4°) and G5 (Φ = 79.8° ± 0.9°, Ψ = 1.7° ± 2.5°) (**Fig. 2L**). This turn reverses the peptide backbone and allows UTag to adopt a β-hairpin conformation. One edge of the resulting UTag framework from residue 6-10 is anchored by a network of backbone-to-backbone hydrogen bonds with the intrabody CDR H3 loop. The interior face of the UTag β-hairpin (**Fig. 2M**) contributes to a hydrophobic core by the burial of peptide residues M1, L3 and W7 against the intrabody. The intrabody side of the pocket is defined by Y105, Y109, and F176. Although the side-chain of Y172 was not fully resolved in the X-ray data, the C_α_ orientation and local environment suggest peripheral participation in this hydrophobic cluster. Another key anchor within the UTag peptide is residue R6, the positively charged guanidinium headgroup of R6 forms hydrogen bonds with the phenolic hydroxyl group (–OH) of anti-UTag-IB Y59 and the backbone carbonyl oxygen (–C=O) of K236 (**Fig. 2N**). This bridging interaction network is conserved across all four copies in the asymmetric unit (PDB: 8V8F) and both copies of the parental CDR-donor Fab (PDB: 2HRP). We also observed polar interactions involving anti-UTag-IB E103 and Y110 that were not present in the parental structure (PDB: 2HRP). In one of the four copies, the hydroxyl group (-OH) of E103 forms a hydrogen bond with the side-chain ammonium group (-NH₃⁺) of UTag K10 (**Fig. 2O**). Meanwhile, in three of the four copies in the asymmetric unit, the phenolic hydroxyl group (–OH) of Y110 forms a hydrogen bond with the side-chain ammonium group (-NH₃⁺) of UTag K8 (**Fig. 2O**). These observations suggest that UTag retains significant flexibility and can adopt multiple productive conformations within the crystal lattice. Despite the variability in certain side-chain contacts, the UTag β-hairpin is preserved. Stabilizing intra-chain contributions include backbone-backbone hydrogen bonds as well as a R6 sidechain-backbone hydrogen bond (**Fig. 2P**).

### Crystal structure reveals the designed cysteine-free anti-UTag intrabody retains native folding conformation

To ensure robust folding of the anti-UTag intrabody in both reducing or oxidizing intracellular environments, we engineered a cysteine-free intrabody variant, anti-UTag-IB(ΔCys). This design incorporated five mutations (L4I, C22A, C96A, C163V, and C232A) to stabilize the protein core in the absence of disulfide bonds. To validate its structural integrity, we determined its crystal structure (PDB: 9N97), which showed remarkable agreement with both the AlphaFold3 (AF3) prediction (RMSD_Cα_ of 0.6 Å; **Fig. 3A**) and the parental anti-UTag-IB structure (PDB: 7URL; RMSD_Cα_ of 0.4 Å; **Fig. 3B)**.

**Figure 3.**
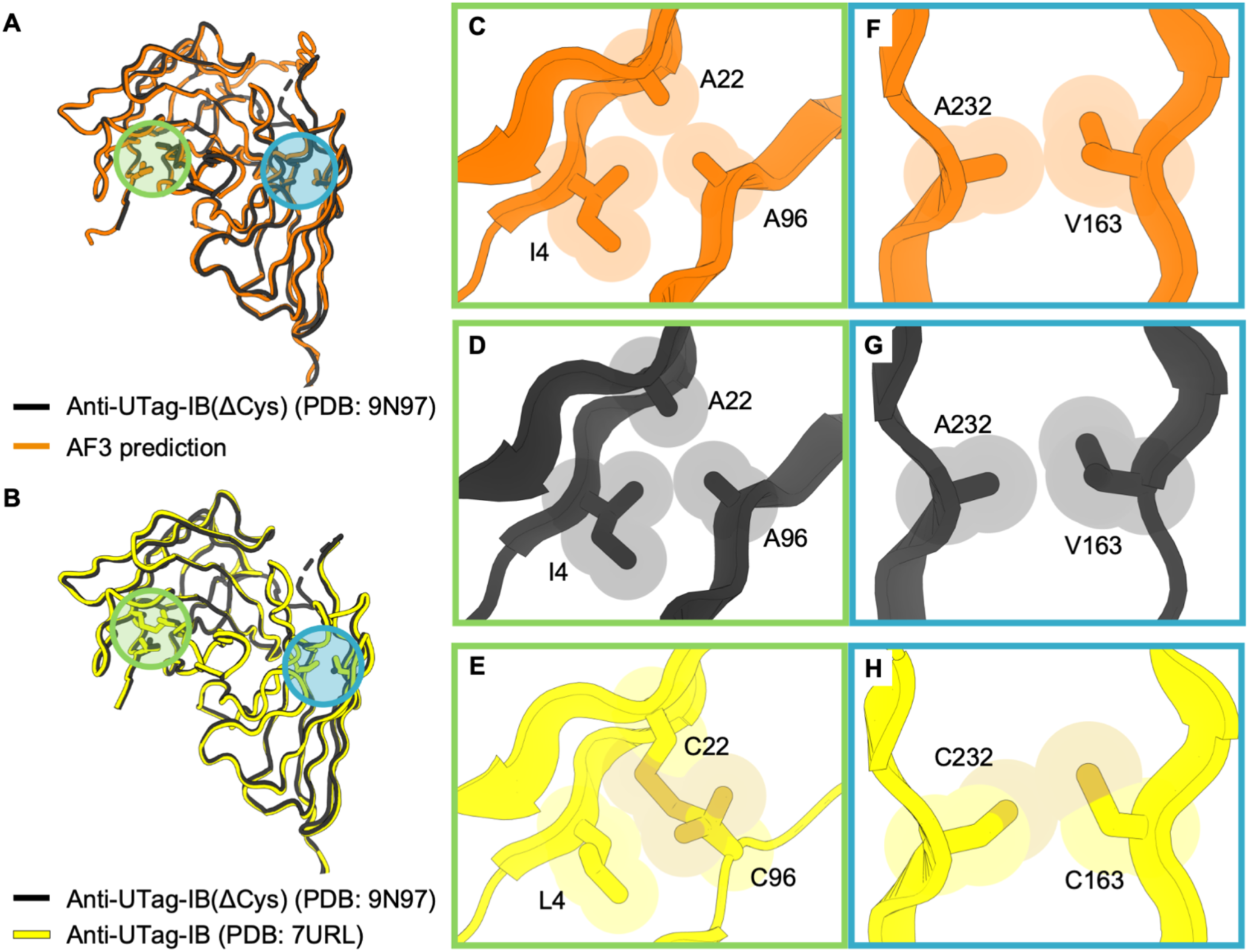
Designed anti-UTag-IB(ΔCys) retains native folding conformation. **(A)** Aligned structures of anti-UTag-IB(ΔCys) with AlphaFold3 prediction, showing a high degree of structural similarity (RMSD_Cα_ = 0.6 Å). **(B)** Aligned structures of anti-UTag-IB(ΔCys) with anti-UTag-IB. RMSD_Cα_ = 0.4 Å. The locations of the cysteine substitutions are highlighted in blue and green boxes in A and B. **(C-E)** Detailed structure views of the highlighted green box in A and B, showing successful elimination of the first disulfide bond via L4I, C22A, and C96A mutations. **(F-H)** Detailed structure views of the highlighted blue box in A and B, showing the second disulfide bond elimination via C163V and C232A substitutions.

Detailed inspection of the first mutation cluster (L4I, C22A, and C96A) confirmed that these mutations successfully eliminated the disulfide bond while preserving a tightly packed hydrophobic core, closely matching both the AF3 model and the parental anti-UTag-IB structure (**Figs. 3C-3E**). Similarly, the second mutation cluster (C163V and C232A) exhibited strong structural concordance (**Figs. 3F-3H**), confirming that the hydrophobic substitutions effectively stabilized the protein interface despite the loss of covalent disulfide bonds.

Global structural comparisons revealed minimal conformational deviations between anti-UTag-IB(ΔCys) (PDB: 9N97) and the parental anti-UTag-IB structure (PDB: 7URL) (**Fig. 3B**). Local analysis of five-residue segments surrounding each mutation site showed subtle RMSD_Cα_ shifts, ranging from 0.05 Å (L4I) to 0.14 Å (C232A) (**Table S2**). Furthermore, anti-UTag-IB(ΔCys) successfully crystallized in the same P2_1_2_1_2_1_ space group as the parental structure 7URL (**Fig. S2A**, **Tables S3-S4**), indicating conserved crystal packing. This preservation of lattice symmetry further supports that the overall protein fold and surface presentation remain essentially unchanged, demonstrating that the disulfide bonds were not critical for maintaining the structural integrity or the crystal arrangement. Collectively, these results validate our design strategy and yield a robustly folded cysteine-free intrabody.

### Anti-UTag intrabodies exhibit tight binding to UTag in live cells

As suitable candidates for live-cell imaging probes, the anti-UTag intrabodies must bind UTag tightly in live cells. To assess this, we characterized the turnover kinetics of both anti-UTag-IB and anti-UTag-IB(ΔCys) against UTag using fluorescence recovery after photobleaching (FRAP) assay in live cells. In parallel, we benchmarked their binding performance against established tagging systems, including SunTag, ALFA-tag, and HA tag. In these FRAP assays, a defined nuclear region was photobleached in cells expressing mEGFP-fused intrabodies, which colocalized in the nucleus with co-expressed 1x epitope-tagged mCherry-H2B (**Figs. 4A-4E**, left). As H2B displays extremely slow turnover, fluorescence recovery in the mEGFP channel primarily reflects the exchange kinetics of intrabodies.^22,33^ Accordingly, longer recovery times indicate tighter binding between intrabodies and their cognate tags.^34^

**Figure 4.**
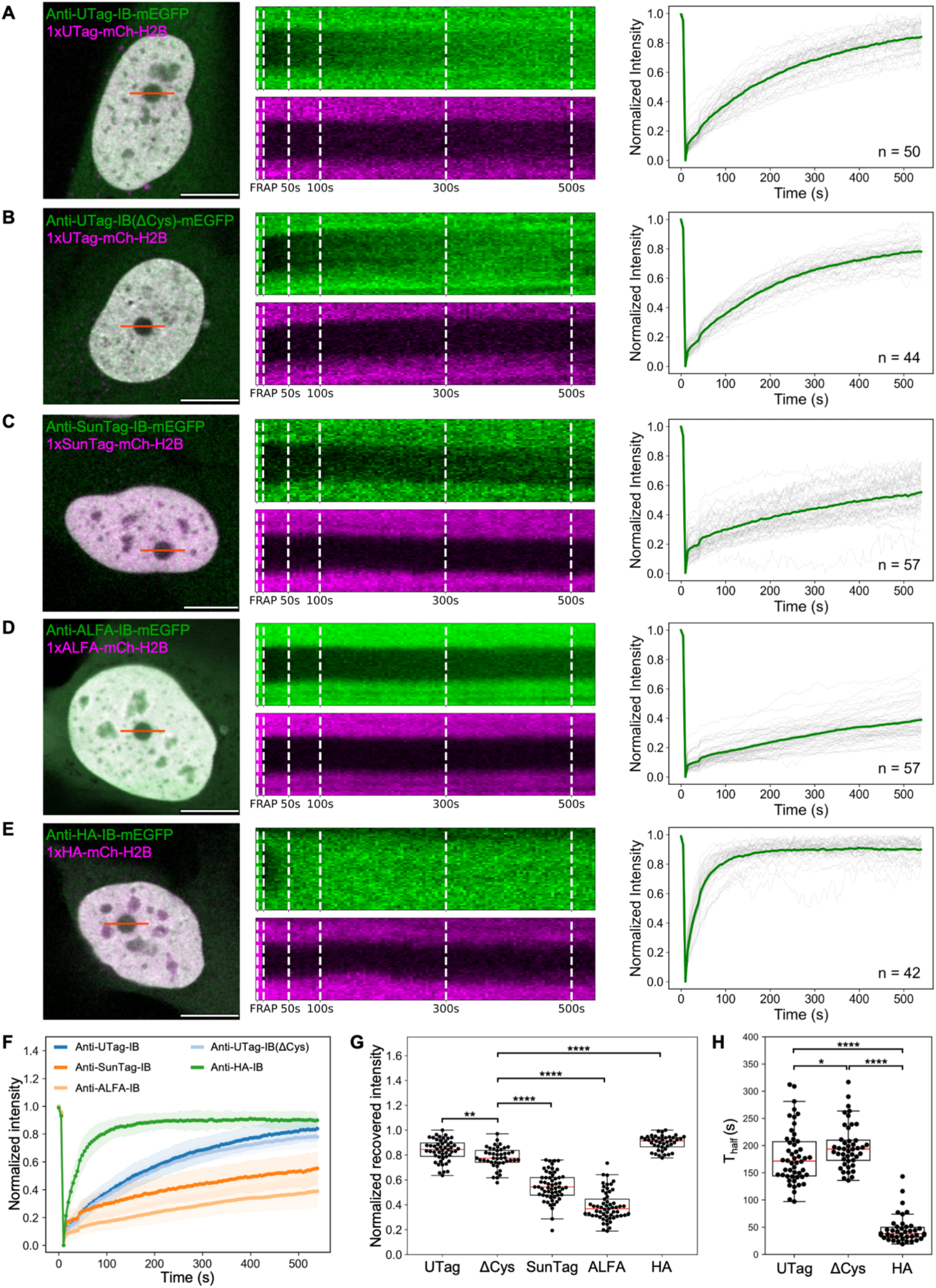
Anti-UTag intrabodies bind UTag tightly in live cells. **(A-E)** FRAP results of each tested tagging system. Left, representative cell images with the photobleached region of interest (ROI). Middle, kymographs of fluorescence recovery over time along transverse lines (red lines; 50 pixels) across the bleached ROIs (green, intrabody-mEGFP; magenta, 1xepitope tag-mCh-H2B). Right, normalized FRAP recovery curves for individual cells (gray) and the mean (green) within the ROI. **(A)** Anti-UTag-IB-mEGFP (n = 50 cells). **(B)** Anti-UTag-IB(ΔCys)-mEGFP (n = 44 cells). **(C)** Anti-SunTag-IB-mEGFP (n = 57 cells). **(D)** Anti-ALFA-IB-mEGFP (n = 57 cells). **(E)** Anti-HA-IB-mEGFP (n = 42 cells). **(F)** Mean FRAP recovery curves for all tagging systems. **(G)** Normalized fluorescence recovery at the endpoint of the FRAP experiment. **(H)** Half-recovery time plots of the FRAP experiment. Mann-Whitney U test. **** p < 0.0001, *** p < 0.001, ** p < 0.01, and * p < 0.05. Only significant differences are indicated. Scale bars: 10 µm. In the box-and-whisker plots, the median is shown in red.

As shown in **Figs. 4A-4E** middle panels, in the FRAP kymograph of each tested tagging system, the magenta channels (1x epitope-tagged-mCh-H2B) showed negligible recovery in the photobleached spots over ∼10 min, consistent with the slow turnover kinetics of H2B. In contrast, the green channel (Intrabody-mEGFP) displayed distinct recovery kinetics across tagging systems (**Figs. 4A-4F**). Consistent with their very high reported affinities (0.38 nM for SunTag^24^ and 26 pM for ALFA-tag^35^), the SunTag and ALFA-tag systems showed limited recovery, reaching only 55% ± 2% (SunTag) and 39% ± 2% (ALFA-tag) of pre-bleach intensity, respectively, at the endpoint of the FRAP experiments (**Fig. 4G**; **Table S6**). In comparison, anti-UTag-IB and anti-UTag-IB(ΔCys) showed substantially higher recovery, reaching 84% ± 1% and 78% ± 1%, respectively, similarly to our previously developed anti-HA-IB (90% ± 1%)^22^ (**Fig. 4G**; **Table S6**).

The mean half-recovery times (t**_1/2_**) were ∼177 s for anti-UTag-IB and ∼194 s for anti-UTag-IB(ΔCys) (p = 0.03), indicating modestly slower turnover and thus slightly tighter binding for the cysteine-free intrabody (**Fig. 4H** and **Table S6**). Consistent with these FRAP results, *in vitro* measurements showed that anti-UTag-IB(ΔCys) exhibited a slightly lower K_D_ than anti-UTag-IB (**Fig. S4**; 11.7 ± 4.8 nM for anti-UTag-IB(ΔCys) vs. 15.0 ± 2.9 nM for anti-UTag-IB). Notably, both anti-UTag intrabodies showed substantially longer half-recovery times than previously developed anti-HA intrabody, which has been widely used for tracking nascent and mature proteins in live cells and organisms^17,22^ (**Fig. 4H** and **Table S6**).

Together, these results indicate that anti-UTag intrabodies bind UTag with high affinity in live cells and are well suited for applications such as single-mRNA translation tracking in live cells, thereby having the potential to expand the currently limited toolbox.

### Anti-UTag intrabodies show superior thermostability

To generate properly folded scFvs in live cells, one prevailing hypothesis is that enhanced *in vitro* thermostability improves their folding performance in live cells.^28^ For example, anti-SunTag intrabody was developed through multiple rounds of engineering to improve its thermostability, ultimately yielding a variant that folds correctly and binds SunTag with high affinity and specificity in live cells.^24,28^ Having successfully engineered two anti-UTag intrabodies that fold properly in live cells from a misfolded wild-type scFv, we sought to characterize their thermostability *in vitro* to assess whether this positive correlation holds for anti-UTag intrabodies development.

To this end, we expressed wild-type anti-UTag-scFv-mEGFP, anti-UTag-IB-mEGFP, and anti-UTag-IB(ΔCys)-mEGFP in *E. coli,* obtained their clarified cell lysates, incubated them at elevated temperatures, centrifuged them to separate soluble and aggregated fractions, then analyzed the soluble fractions by western blot (**Fig. 5A**). The results (**Figs. 5B-5C, S5A**) show that the wild-type anti-UTag-scFv was largely aggregated at 50 °C, with undetectable soluble protein remaining. In contrast, anti-UTag-IB displayed markedly improved thermostability, retaining full solubility at 50 °C and ∼75% solubility at 60 °C, but losing solubility at 70 °C (**Figs. 5B-5C, S5B**), indicating that the scaffold-donor 15F11 enhanced thermostability, thereby facilitating proper folding in live cells. Excitingly, the cysteine-free variant, anti-UTag-IB(ΔCys), retained ∼40% solubility even at 70 °C (**Figs. 5B-5C, S5C**), indicating a further improvement in thermostability upon cysteine removal.

**Figure 5.**
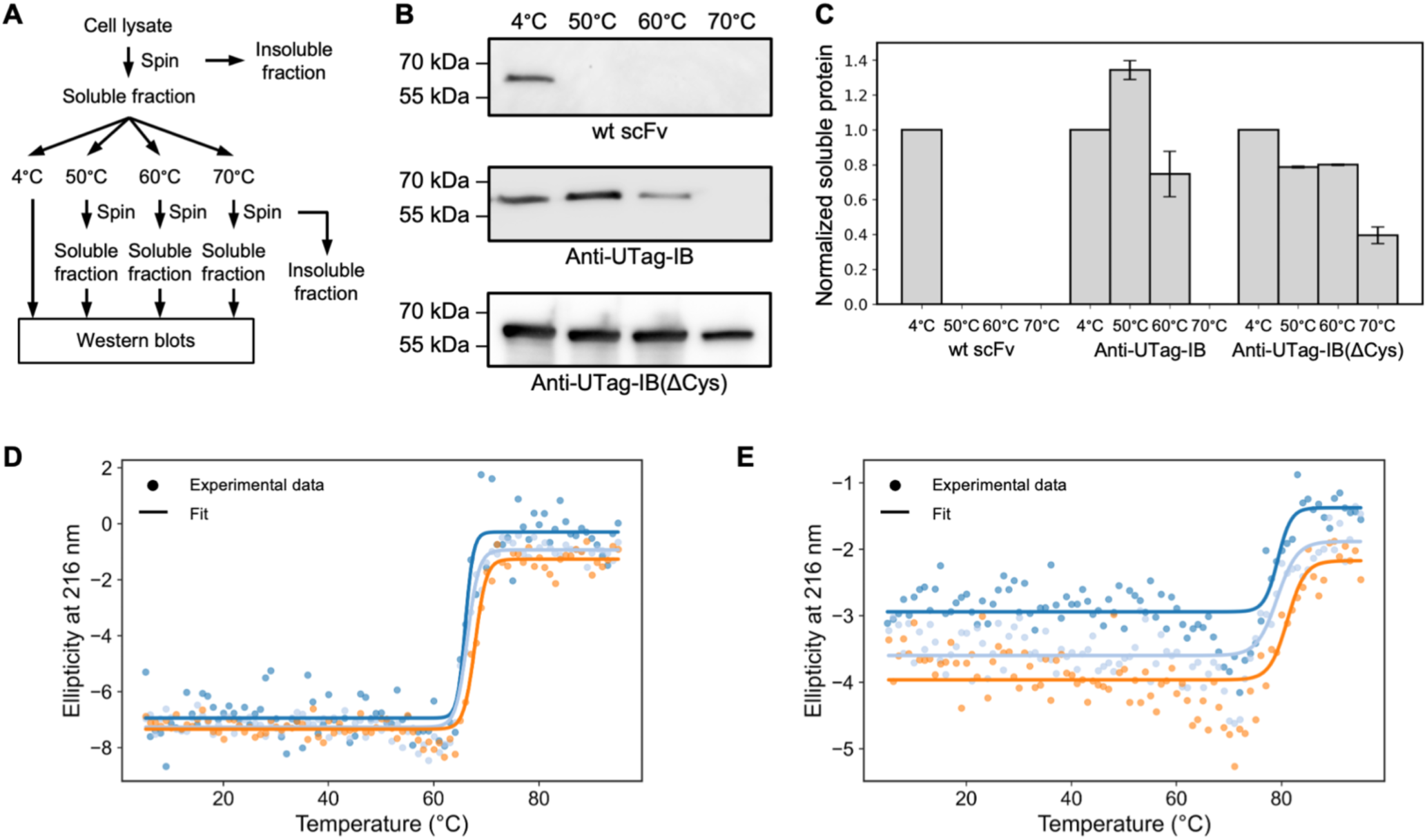
Anti-UTag intrabodies show improved thermostability. **(A)** Schematic illustrating the procedure used to assess intrabody thermostability. **(B)** Representative western blots of soluble fractions after incubation for 15 min at 4 °C, 50 °C, 60 °C, and 70 °C for wild-type anti-UTag-scFv (top), anti-UTag-IB (middle), and anti-UTag-IB(ΔCys) (bottom). **(C)** Quantification of western blots in (B). Error bars represent SEM from two independent experiments. **(D)** Circular dichroism (CD) thermal denaturation of anti-UTag-IB. **(E)** CD thermal denaturation of anti-UTag-IB(ΔCys). CD curves shown in different colors in (D) and (E) represent three independent experiments.

To validate these thermostability results, we measured the melting temperatures (T_m_) of purified anti-UTag-IB and anti-UTag-IB(ΔCys). Anti-UTag-IB exhibited a T_m_ of 66.76 ± 0.46 °C (**Fig. 5D**), whereas anti-UTag-IB(ΔCys) showed an increased T_m_ of 79.87 ± 0.52 °C (**Fig. 5E**). These results are consistent with the thermostability results and confirm that cysteine removal enhances thermostability. Collectively, these findings support the hypothesis that improved thermostability *in vitro* promotes folding *in vivo*.

### An optimized linker enables efficient intrabody binding to tandem UTags without perturbing reporter cellular localization

For live-cell imaging applications, epitope tags are often repeated multiple times to enhance signal-to-noise. For example, tracking single mRNA translation in live cells typically requires tandem arrays of 24x tags to enable long-term imaging. To extend this strategy to anti-UTag intrabodies, we optimized the linker length between tandem UTags to maximize intrabody binding stoichiometry.

We designed three linker lengths: 5-, 9-, and 13-aa and evaluated their performance in cells co-expressing a mitochondrial outer membrane protein mitoNEET^36^ (Mito) fused to mCh-4xUTag (Mito-mCh-4xUTag) and anti-UTag-IB-mEGFP (**Fig. 6A**). Imaging was performed under identical conditions alongside cells expressing the control construct mEGFP-mCh-H2B (**Fig. 6A**). Fluorescence intensities on mitochondria were quantified in both mEGFP and mCh channels, and the mEGFP/mCh ratios were normalized to the control. As shown in **Fig. 6B**, relative to the control construct (1.01 ± 0.16), the 5-aa linker increased the ratio to 3.47 ± 1.18 (p < 0.0001). Similar increases were observed for the 9-aa (3.25 ± 0.87; p < 0.0001) and 13-aa linkers (3.20 ± 1.18; p < 0.0001). No significant differences were detected among the linker lengths, indicating that the 5-aa linker provides sufficient spacing for maximal intrabody binding (**Fig. 6B**). Accordingly, we selected the 5-aa linker for constructing 24xUTag arrays for single-mRNA translation tracking in live cells.

**Figure 6.**
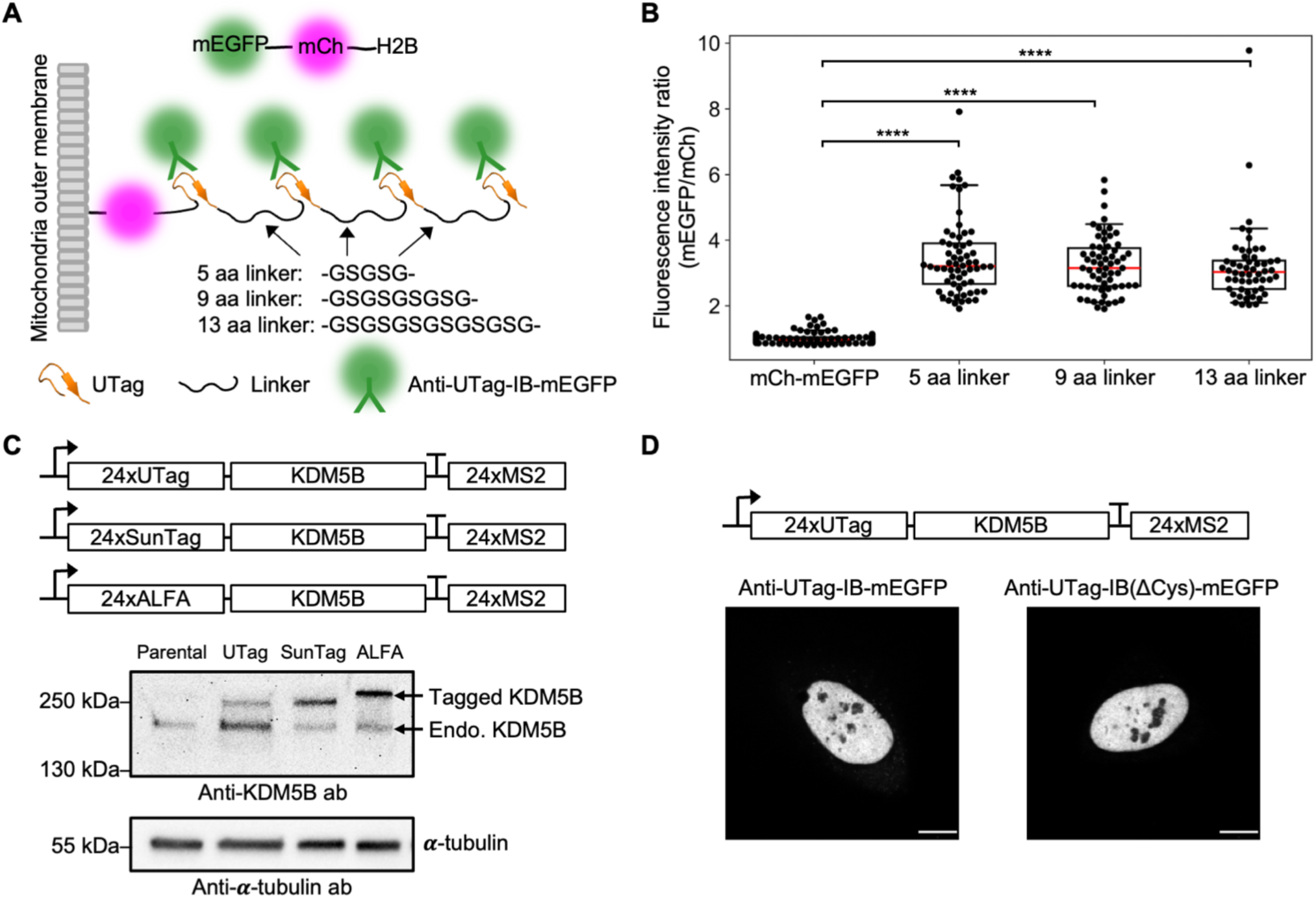
An optimized linker enables efficient intrabody binding to tandem UTags without perturbing reporter cellular localization. **(A)** Schematic illustrating three tested linker lengths (5, 9, and 13 aa) designed to enable efficient binding of anti-UTag-IB-mEGFP to 4xUTag fused to Mito-mCh (Mito-mCh-4xUTag). The control construct mEGFP-mCh-H2B is shown at the top. **(B)** Box-and-whisker plot of the mEGFP/mCh fluorescence intensity ratio, normalized to the ratio of mEGFP-mCh-H2B construct. From left to right, n = 128, 62, 62, 54 cells respectively. **(C)** Representative western blot of 24xUTag-, 24xSunTag-, and 24xALFA-tagged KDM5B reporters, confirming expression of full-length proteins. **(D)** Representative cell images showing nuclear localization of 24xUTag-KDM5B visualized by anti-UTag-IB-mEGFP and anti-UTag-IB(ΔCys)-mEGFP. Mann-Whitney U test. **** p < 0.0001. Only significant differences are indicated. Scale bars: 10 µm. In the box-and-whisker plots, the median is shown in red.

We next fused the 24xUTag to the N-terminus of a previously studied reporter protein KDM5B^2^ and assessed whether full-length protein expression could be achieved, alongside constructs 24xSunTag- or 24xALFA-tagged KDM5B (**Fig. 6C**, top). After 12 h of expression, KDM5B in cell lysate was analyzed by western blot. As shown in **Figs. 6C** and **S6**, all tagged KDM5B constructs displayed a single band at the expected molecular weight, indicating successful expression of full-length proteins.

Finally, we tested whether intrabody binding to 24xUTag-KDM5B perturbs the nuclear localization of KDM5B. Cells were imaged 24 h after transfection with either anti-UTag-IB-mEGFP/24xUTag-KDM5B or anti-UTag-IB(ΔCys)-mEGFP/24xUTag-KDM5B constructs. As shown in Figs**. 6D**, **S7A**, and **S7B**, both intrabodies exhibited robust nuclear localization of KDM5B, indicating that the binding between the intrabodies and 24xUTag array does not interfere with reporter protein localization and is therefore unlikely to disrupt its function. In contrast, cells expressing anti-SunTag-IB-mEGFP/24xSunTag-KDM5B showed weaker nuclear localization (**Fig. S7C**). Notably, cells expressing anti-ALFA-IB-mEGFP/24xALFA-KDM5B exhibited almost no nuclear localization (**Fig. S7D**), likely due to the extremely tight binding of the anti-ALFA-IB for the ALFA-tag. Taken together, these results demonstrate that the 24xUTag system, in combination with either anti-UTag intrabody, provides an effective and minimally perturbative tagging strategy for live-cell imaging.

### Benchmarking anti-UTag intrabodies for single-mRNA translation tracking in live cells against established tagging systems

To expand the existing limited toolbox for live-cell single-mRNA translation tracking, we demonstrate here that our developed anti-UTag-IB and anti-UTag-IB(ΔCys) enable robust translation imaging in live cells and benchmark their performance against established tagging systems.

As UTag is a structured tag, we adopted a similar tandem strategy as used for SunTag and ALFA-tag to generate a 24xUTag array. For systematic comparison, 24xUTag, 24xSunTag, or 24xALFA were fused to the N-terminus of the same reporter protein KDM5B (**Fig. 7A**). For imaging, all intrabodies were constructed with identical C-terminal fusions of mEGFP and the solubilization domain GB1. These intrabodies specifically light up translation spots by binding their cognate tags as they emerge from actively translating ribosomes (**Fig. 7B**). Translation spots were tethered to the plasma membrane through interactions between MS2 stem-loops in the reporter 3’ untranslated region (UTR) and MS2 coat protein (MCP) fused to a membrane targeting domain CAAX, as applied previously (**Fig. 7B**).^5^ We observed diffraction-limited translation spots that could be tracked over time across all four tested tagging systems (**Movies S1-S4**). These spots were confirmed as *bona fide* active translation spots by puromycin treatment. Quantitative analysis revealed comparable spot intensities across all tagging systems (**Fig. 7C**). Similarly, signal-to-noise ratios (SNRs) were comparable across systems (**Fig. 7D**). The only notable optical difference was a slightly larger spot size for the SunTag (0.36 µm vs. 0.33 µm for the others) (**Fig. 7 E**). This difference may reflect the longer length of SunTag (19-aa) relative to UTag (11-aa) and ALFA-tag (13-aa). Overall, these results demonstrate that all four tagging systems produce robust, high-contrast single-mRNA translation signals with comparable quality, enabling quantitative analysis.

**Figure 7.**
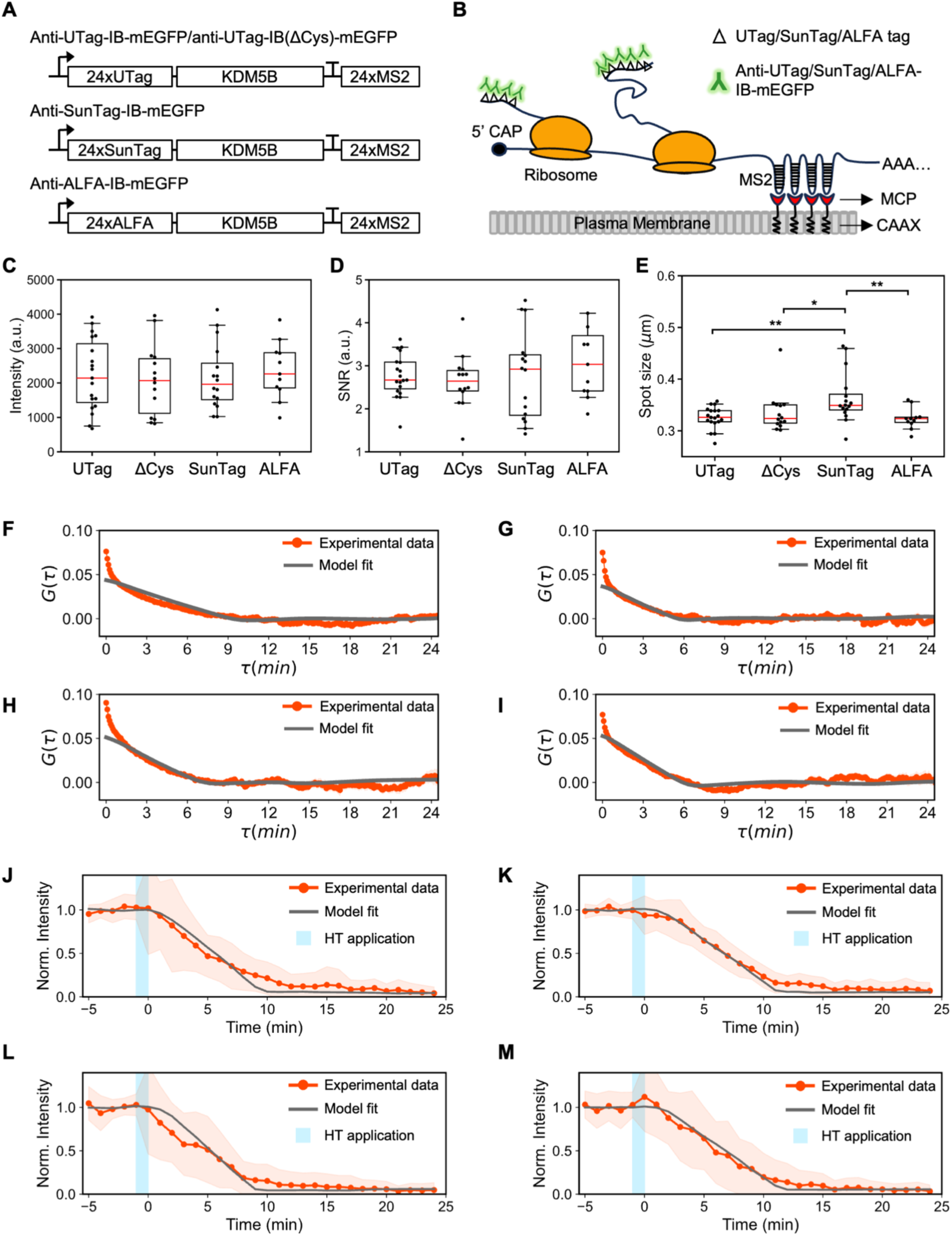
Benchmarking anti-UTag intrabodies for single-mRNA translation tracking in live cells against established tagging systems. **(A-B)** Schematic illustrating single-mRNA translation tracking using four tagging systems in live cells. **(C-E)** Quantitative analysis of translation spots visualized by the four tagging systems: (C) spot intensity calculated using the background subtraction method; (D) signal-to-noise ratio (SNR); (E) spot size determined by 2D Gaussian fitting (full width at half maximum (FWHM)). **(F-I)** Autocorrelation functions (ACFs) calculated from single-spot fluorescence trajectories for the four tagging systems: (F) Anti-UTag-IB-mEGFP (n = 19 cells, 302 trajectories); (G) Anti-UTag-IB(ΔCys)-mEGFP (n = 14 cells, 166 trajectories); (H) Anti-SunTag-IB-mEGFP (n = 16 cells, 136 trajectories); (I) Anti-ALFA-IB-mEGFP (n = 11 cells, 157 trajectories). Red lines represent experimental data; gray lines show TASEP model fits. **(J-M)** Harringtonine (HT) run-off assays measuring translation elongation rates for the four tagging systems, showing the time course of translation spot intensity decay after harringtonine treatment: (J) Anti-UTag-IB-mEGFP (n = 32 cells); (K) Anti-UTag-IB(ΔCys)-mEGFP (n = 22 cells); (L) Anti-SunTag-IB-mEGFP (n = 33 cells); (M) Anti-ALFA-IB-mEGFP (n = 32 cells). Red lines represent experimental data; gray lines show TASEP model fits. Mann-Whitney U test. ** p < 0.01 and * p < 0.05. Only significant differences are indicated. In the box-and-whisker plots, the median is shown in red.

To further compare translation kinetics measured using the UTag system to others, we applied autocorrelation function (ACF) analysis to the intensity trajectories of translation spots, an approach previously used to study single-mRNA translation kinetics.^2,37^ Across all four tagging systems, ACF analysis exhibited similar decorrelation times, ranging from approximately 6 to 9 min (**Figs. 7F–7I**, red lines).

To extract translation kinetics from the ACF analysis, we employed a totally asymmetric exclusion process (TASEP) model in which ribosomes initiate at a rate *k*_*i*_ (s⁻¹), elongate at a rate *k*_*e*_ (aa/s), and are subject to steric exclusion with a ∼9-codon ribosome footprint (**Fig. S8A**). Fluorescence intensity of translation spots was modeled as the cumulative signal from exposed tags within the N-terminal 24xtag array on nascent chains (**Fig. S8B**). We first validated the model by recovering ground-truth parameters from simulated data (**Figs. S8C-S8D**), then fitted simulated ACFs to experimental ACFs to infer *k*_*i*_ and *k*_*e*_for each tagging system (**Figs. 7F-7I; Figs. S9A-S9D**). The best-fit model revealed *k*_*i*_ranging from 0.031-0.063 s⁻¹ (approximately 2-4 initiation events per minute) and *k*_*e*_of 3.1-5.3 aa/s across all four tagging systems (**Table S7**). These values correspond to initiation intervals of 16-32 s and are consistent with reported mammalian translation kinetics^37^. The model further predicts ribosome densities of 7.1-10.7% and an average of 17-24 ribosomes per mRNA at steady state, which are consistent with recent measurements (<12%) reported by Lamberti *et al.*^38^.

To validate the translation kinetics obtained by ACF analysis, we applied an orthogonal harringtonine run-off assay to measure the elongation rates across all four tagging systems. Harringtonine inhibits translation initiation by binding to the 60S ribosomal subunit and preventing ribosome assembly. Upon treatment, elongating ribosomes continue to translate along the mRNA, while no new ribosomes are loaded, resulting in a progressive decay of fluorescence intensity at translation spots. To obtain translation elongation rates, we measured the run-off time required for translation spot intensities to decrease to basal levels. In some cases, translation spots exhibited no response to harringtonine treatment, maintaining stable intensities throughout the experiment. Such harringtonine-resistant spots have been reported and may correspond to stalled ribosomes. To avoid biasing the analysis, these non-responsive spots were excluded. The resulting intensity time courses (**Figs. 7J-7M**, red lines) represent the normalized mean intensity of responsive translation spots across all imaged cells for each tagging system.

To extract elongation rates from the harringtonine data, we fixed the initiation rate at 0.03 s⁻¹ (representative value from the ACF analysis) and performed a grid search over 15 evenly spaced elongation rates spanning 1.0-8.0 aa/s. For each candidate elongation rate, model-data agreement was quantified by chi-squared fitting over the harringtonine post-treatment window. The best-fit curve for each tagging system is shown in **Figs. 7J-7M** (gray lines) and the corresponding elongation rates are summarized in **Table 1**. Overall, the elongation rates derived from the harringtonine run-off assay were comparable with those obtained from ACF analysis (**Table 1**).

**Table 1.** Elongation rate (*k_e_*) comparison from ACF analysis and harringtonine (HT) run-off assay.

| $k_e$ (aa/s) | Anti-UTag-IB | Anti-UTag-IB(ΔCys) | Anti-SunTag-IB | Anti-ALFA-IB |
| --- | --- | --- | --- | --- |
| ACF | 3.11 | 5.29 | 4.33 | 4.84 |
| HT | 3.50 | 3.00 | 4.00 | 3.00 |

Taken together, the translation kinetics measured with all four tagging systems are largely comparable in either ACF analysis or harringtonine run-off assay, demonstrating that our newly developed anti-UTag-IB and anti-UTag-IB(ΔCys) provide robust and complementary tools for studying translation kinetics in live cells.

## Discussion

In this study, we used our previously identified stable scFv scaffold, 15F11, to mine antibody fragments in the PDB and identified an anti-UTag Fab with high sequence identity (∼85%). Grafting its CDR loops onto the 15F11 scaffold generated a novel and tight binding anti-UTag intrabody that folds correctly in live cells and enables real-time tracking of single-mRNA translation *in vivo*. In addition to our previously developed anti-HA and anti-FLAG intrabodies^22,23^, we have now successfully generated three intrabodies using the same loop-grafting strategy, the 15F11 scaffold donor, and a sequence-guided approach for selecting matched CDR loops. Notably, this engineering process requires only a single loop grafting step without extensive screening, highlighting its potential to accelerate intrabody development as antibody sequence data continue to expand.

A leading hypothesis in the development of scFv-based intrabodies is that reducing their folding dependence on disulfide-bond formation in the scaffold improves intracellular functionality. In live cells, disulfide bonds are typically not formed efficiently in the reducing environment of the cytoplasm, where intrabodies are synthesized and folded. However, this hypothesis has not been rigorously validated experimentally. In this study, our solved crystal structure of the anti-UTag intrabody indicates a markedly reduced propensity for disulfide bond formation (0% and 8%), suggesting that the intrabody folds largely independently of disulfide bonds. To determine whether this reduced dependence is an intrinsic property of the scaffold donor 15F11, we re-refined its previously reported crystal structure (PDB: 5B3N) and found a similarly significantly reduced propensity for disulfide bond formation (0% and 60%). These results indicate that the folding of the 15F11 scaffold itself exhibits substantially reduced reliance on disulfide-bonds, and that grafting the anti-UTag Fab CDR loops further diminishes this dependence. Consistent with these structural findings, the anti-UTag intrabody is strongly enriched in the cell nucleus with negligible cytoplasmic signal when co-expressed with 4xUTag-mCh-H2B (**Figs. 1D-1F**). Moreover, we observed no substantial aggregation of the anti-UTag intrabody in live cells, further supporting its proper and robust folding.

Although the folding of the anti-UTag intrabody exhibits minimal dependence on disulfide-bonds and has been successfully used to track UTag-tagged proteins in both the cytoplasm and nucleus, the four cysteine residues in its scaffold could potentially form non-native disulfide bonds in other intracellular compartments, such as the ER lumen, where an oxidizing environment is maintained. To address this limitation, we further engineered the scaffold to completely eliminate all cysteine residues. The resulting cysteine-free anti-UTag intrabody displays increased thermostability (T_m_ is ∼80 °C) while maintaining its binding affinity to UTag both *in vivo* and *in vitro*. We anticipate that this cysteine-free variant will enable studies of translation within the ER lumen, a process that remains largely unobservable due to the current lack of suitable tools.

Our developed anti-UTag intrabodies expand the currently available toolbox for live-cell translation tracking. We compared their performance with other widely used tagging systems, including 24xSunTag and 24xALFA, using the same nuclear protein KDM5B as a reporter. We observed robust nuclear localization of 24xUTag-KDM5B, as visualized by both anti-UTag intrabodies, while 24xALFA-KDM5B was largely depleted from the nucleus, likely due to the super tight interaction between the ALFA-tag and the anti-ALFA nanobody (26 pM)^35^. The nuclear localization of 24xSunTag-KDM5B was intermediate, which may result from high expression of the anti-SunTag intrabody. These results indicate that excessively high binding affinity can be detrimental, as it can interfere with proper intracellular localization of the reporter protein. In contrast, anti-UTag intrabodies exhibit a favorable binding affinity to UTag that supports both efficient translation tracking and correct localization of the mature protein.

We further compared translation kinetics measured using 24xUTag-KDM5B, 24xSunTag-KDM5B, and 24xALFA-KDM5B to assess whether different tagging systems influence translation kinetics of the reporter. Two complementary approaches were employed: ACF analysis and harringtonine run-off assays. ACF analysis determined elongation rates ranging from 3.1-5.3 aa/s across the different tagging systems, whereas harringtonine run-off assays yielded slightly slower but comparable rates of 3.0-4.0 aa/s. This 1-2 aa/s discrepancy is consistent with previous observations that different approaches can produce different elongation rates. For example, Pichon *et al*. reported 13.2 aa/s from harringtonine run-off assays versus 18 aa/s from FRAP using the same 56xSunTag-Ki67 reporter.^3^ Several factors may contribute to these differences. Harringtonine run-off assays are sensitive to drug pharmacokinetics, including batch-to-batch variability in drug potency and the precise timing of initiation inhibition. In contrast, ACF analysis relies on accurate particle tracking and fluorescence intensity quantification over long timescales. Tracking errors, photobleaching artifacts, and mRNA diffusion in and out of the focal plane (the diffusion still occurs when tethered to cell plasma membranes) can also introduce noise that affects correlation decay. Given these complementary strengths and limitations, we consider both approaches to bracket the true elongation rate. The consistency of our measurements across multiple tagging systems, together with their agreement with reported values in the literature (2-10 aa/s), supports the robustness of our analysis.^37^ Importantly, these results demonstrate that the UTag tagging system performs comparably to established and time-tested tagging systems, indicating that it can be readily combined with SunTag and ALFA-tag systems for multiplexed imaging applications.

Our TASEP-based model successfully captured the long-timescale decay of the experimental ACF analysis but underfitted the rapid initial decay at short time lags (**Figs**. **7F-7I**). This discrepancy persisted despite extensive parameter optimization within biologically plausible ranges, suggesting a structural limitation of the model rather than an issue with parameter estimation. To validate our implementation, we simulated traces using known parameters and accurately recovered them (**Figs**. **S8C-S8D**), confirming the internal consistency of the model. The fast initial decay may arise from several factors. First, as noted by Coulon *et al*.^39^, the zero-lag region of ACF analysis is particularly sensitive to shot noise. Second, rapid mRNA diffusion could cause intensity fluctuations on timescales shorter than the ribosomal elongation cycle. Third, the binding/unbinding kinetics of intrabodies could contribute to the fluorescence fluctuation during the initial decay. Fourth, a key limitation of our model is the assumption of constant-rate (Poisson) initiation instead of bursting initiation. In bursting initiation, ribosomes load in clusters followed by inactive periods, which could generate the observed rapid decorrelation at short time lags. Future work should extend the model to explicitly incorporate bursting initiation using a multi-state initiation model.

## Methods

### Plasmids

All plasmids and their corresponding sequences used in this study are listed in **Table S8**. Plasmid sequences were verified by whole-plasmid nanopore sequencing (Quintara Biosciences). Plasmids used for imaging experiments were prepared using the ZymoPURE II Plasmid Midiprep Kit (Zymo Research) and adjusted to a final concentration of ∼100 ng/µL.

### U-2 OS cell culture

U-2 OS cells (ATCC HTB-96) were maintained in DMEM medium (Gibco) supplemented with 10% (v/v) fetal bovine serum (FBS; Gibco), 1 mM L-glutamine (Gibco) and 1% (v/v) penicillin-streptomycin (Gibco) in a humidified incubator at 37 °C with 5% CO_2_.

### Transfection

Plasmids prepared using the ZymoPURE II Plasmid Midiprep Kit (Zymo Research) were diluted in ultrapure water to a final concentration of ∼100 ng/µL. Prior to transfection, U-2 OS cells seeded in 35 mm MatTek imaging dishes or standard tissue culture dishes were washed twice with 2 mL PBS (pH 7.4). Cells were then supplemented with 1.8 mL prewarmed Opti-MEM (Gibco) and 0.2 mL FBS (final concentration 10%; Gibco) and maintained at 37 °C in a humidified incubator with 5% CO₂ until transfection. For transfection, plasmid DNA (1.25 µg or 2.5 µg) was mixed with Opti-MEM and Lipofectamine LTX reagent (Thermo Scientific), while PLUS reagent was diluted separately in Opti-MEM according to the manufacturer’s instructions. The LTX and PLUS mixtures were then combined and incubated for 5 min at room temperature. The resulting transfection complexes were added dropwise to the cells. The transfection reagent was removed after 3 h and phenol-red-free DMEM medium (Gibco) supplemented with 10% (v/v) fetal bovine serum (Gibco), 1 mM L-glutamine (Gibco) and 1% (v/v) penicillin-streptomycin (Gibco) was added to the cells. The cells were incubated in a humidified incubator at 37 °C with 5% CO_2_ until ready for imaging or harvesting.

### Live-cell imaging conditions

Live-cell images were acquired using a Leica Stellaris 5 confocal microscope equipped with a 63x oil-immersion objective (excluding Fig. 1). During imaging, cells were maintained in a stage-top incubator at 37 °C with 5% CO₂ and 51% relative humidity. Images were collected with a field of view of 512 x 512 pixels at 16-bit depth. The resulting pixel size was 129.89 nm. For z-stack acquisition, images were collected with a step size of 0.3 µm. All image acquisition was performed using Leica Application Suite X (LAS X, version 4.5.0.25531).

Live-cell imaging (excluding FRAP experiments) was performed using resonance scanning mode (8 kHz) with bidirectional scanning and a line averaging of 16. The bidirectional phase offset was set to −87.74, and images were acquired at a zoom factor of 2.78. In Fig. 7 harringtonine run-off experiments, the 488 nm laser was set at 1-2%. In Fig. 7 autocorrelation function analysis experiments, the 488 nm laser was set at 1.5%. In Fig. 6B linker optimization experiments, the 488 nm laser was set at 0.5-2% and the 561 nm laser was set at 1-2%. In Fig. 6D KDM5B localization experiments, the 488 nm laser was set at 1-5%.

For FRAP imaging, time-lapse movies were acquired in FRAP mode using the following parameters: scan speed of 800 Hz, zoom factor of 2.45, and line averaging of 1. Signal detection was performed using a HyD S detector in photon-counting mode. Excitation was achieved with a 488 nm laser at 1.5% power and a 561 nm laser at 2% power. Photobleaching was performed using the 488 nm laser at 20% power with the Zoom-in bleaching mode. The bleached region of interest (ROI) was defined as a circular area with a diameter of 3 µm. Each FRAP acquisition consisted of 10 pre-bleach frames collected at 1 s intervals, followed by 30 post-bleach frames at 1 s intervals and an additional 100 post-bleach frames at 5 s intervals.

Co-localization images shown in Fig. 1 were acquired using an Olympus IX81 spinning-disk confocal microscope (CSU22 head) equipped with a 100x oil-immersion objective (NA 1.40). Sequential imaging was performed using 488 nm and 561 nm lasers for 20 time points. The spinning disk was operated at 1x spin rate, and exposure times were adjusted based on cell brightness. Images were captured using a Photometrics Cascade II CCD camera controlled by SlideBook software (Intelligent Imaging Innovations). Final images displayed in figures were generated by averaging all time points using ImageJ.

### Protein expression and purification of intrabodies

BL21-CodonPlus(DE3)-RIPL cells (Agilent) harboring either pET23b-anti-UTag-IB-TEV-mEGFP or pET23b-anti-UTag-IB(ΔCys)-TEV-mEGFP were grown in 2xYT medium supplemented with Ampicillin (100 mg/L) and Chloramphenicol (25 mg/L) at 37 °C with continuous shaking until reaching an optical density at 600 nm (OD600) of 0.6. Protein expression was induced with 0.4 mM isopropyl β-D-1-thiogalactopyranoside (IPTG; RPI), followed by incubation at 18°C overnight with continuous shaking. Cells were harvested by centrifugation and resuspended in PBS buffer containing 150 mM NaCl and 0.2 mM AEBSF protease inhibitor. Cell lysis was performed by sonication, and the lysate was cleared by centrifugation. The intrabody fusion protein was purified from the clarified lysate using Ni Sepharose Excel affinity chromatography, followed by a linear gradient of 0-500 mM imidazole elution. The eluted intrabody protein was dialyzed against PBS buffer with 500 mM NaCl and 0.2 mM AEBSF to remove imidazole, followed by overnight incubation with house-made TEV protease at a 1:25 enzyme:substrate ratio to cleave the mEGFP tag and 6xHis tag. After the cleavage reaction was completed, the TEV protease was removed by incubating with Ni Sepharose Excel resins (Cytiva) at 4 °C for 4 h or until the supernatant was undetectable for GFP fluorescence. The flowthrough and washes were collected, combined, and concentrated using a Vivaspin 10 kDa MWCO centrifugal filter unit (Sartorius). The concentrated intrabody protein underwent a final purification by size-exclusion chromatography on a HiLoad Superdex 200 PG column equilibrated in HEPES buffer (25 mM HEPES pH 7.9, 12.5 mM MgCl₂, 100 mM KCl, 0.1 mM EDTA, 0.01% NP40, 10% glycerol, and 1 mM DTT). The fractions containing the intrabody were collected, concentrated, and stored at −80 °C after flash freezing in liquid nitrogen.

### Crystallization

Crystallization experiments were performed using the sitting drop vapor diffusion method at 20°C (295K). Initial screening of ∼72 conditions was carried out using three 24-well CrysChem plates, varying pH (5.0-8.0), ammonium sulfate concentration (0.1-4.0 M), and PEG (3350 and 1000) concentrations. The protein-to-precipitant ratio was further optimized (1:2, 1:4, 1:6, and 1:8), with a 1:2 ratio yielding the most favorable crystal growth and size. Optimal crystallization conditions for both wild-type and cysteine-free anti-UTag intrabodies were identified as 2 M ammonium sulfate, 0.1 M BIS-TRIS pH 7.0, and 3% (w/v) PEG 3350. For both intrabodies, crystallization drops were prepared by mixing 0.6 µL of protein solution (15 mg/mL) with 1.2 µL of reservoir solution, corresponding to a 1:2 protein-to-precipitant ratio. Crystals formed overnight under these conditions. For co-crystallization, anti-UTag intrabody was incubated with synthetic UTag peptide (MSLPGRWKPKM; GenScript) at a 10-fold molar excess prior to setup. Co-crystals were obtained using a condition containing 0.2 M ammonium sulfate, 0.1 M BIS-TRIS pH 6.5, and 25% (w/v) PEG 3350 (Hampton Research HR2-134, condition F7). Sitting drops were prepared by mixing 0.6 µL of the protein–peptide complex with 1.2 µL of reservoir solution.

### X-ray diffraction and data processing

In all cases, individual crystals were briefly swished through a cryoprotectant solution prior to flash-freezing in liquid nitrogen. The cryoprotectant consisted of 25% (w/v) PEG 3350 for the wild-type intrabody, 30% (w/v) PEG 3350 with 0.1 M BIS-TRIS pH 7.0 for the cysteine-free variant, and 0.1 M BIS-TRIS pH 5.5, 45% (w/v) PEG 3350, and 1.25 mg/mL UTag peptide for the co-crystals. X-ray diffraction data were collected at beamline 4.2.2 at the Advanced Light Source (ALS). Data were processed using XDS^40^. The wild-type intrabody structure was determined by molecular replacement (MR) using an AlphaFold2-predicted structure as the search model. Model refinement was performed in PHENIX^41^ and COOT^42^, using σ-weighted 2mFₒ–DFc and mFₒ–DFc electron density maps. The resulting wild-type model (PDB: 7URL) was subsequently used as the starting MR model for both the cysteine-free and co-crystal structures, which were refined using the same workflow.

In addition, the previously deposited structure PDB: 5B3N was re-refined to resolve clear mFₒ–DFc difference density (**Fig. S1**, red/green blobs) observed in the regions surrounding the disulfide bonds in the original model. This re-refinement was performed using the same software suite and refinement parameters described above, with coordinates and structure factors obtained from the RCSB PDB, ensuring a consistent baseline for structural comparison.

### Computational design of a cysteine-free anti-UTag intrabody

The cysteine-free anti-UTag intrabody was engineered using a targeted computational strategy focused on preserving structural integrity through maximal heavy-atom conservation and optimized hydrophobic packing. All computational modeling and design were performed using *Rosetta* version 2023.45 (release a6d9ba8). Molecular visualizations, residue neighbor analysis, and initial structure generation were carried out using PyMOL Molecular Graphics System, Version 3.1.3 (Schrödinger, LLC).

The crystal structure of the scaffold donor 15F11 (PDB: 5B3N) served as the starting template. To minimize backbone perturbations, our design protocol followed five core principles: (1) minimizing the total number of mutations, (2) maintaining approximately constant heavy-atom count, (3) avoiding residues with distinct backbone conformational propensities (e.g., Gly and Pro), (4) eliminating unpaired cysteines, and (5) replacing the native disulfide bond volume with hydrophobic side chains. Using PyMOL, we identified neighboring Leu, Ile, and Met residues (hydrophobic side chains containing four heavy atoms) within 4 Å of the native disulfide-forming side chains (specifically the β-carbon and γ-sulfur atoms) for two sites: Group 1 (C24-C98) and Group 2 (C161-C230).

A combinatorial search space was constructed for each site, sampling Ala and Val at former cysteine positions and the [Leu, Ile, Met] set at the identified neighboring sites. To strictly maintain or reduce the total heavy-atom count relative to the native disulfide bond (4 heavy atoms), combinations resulting in a double valine (VV) substitution (6 heavy atoms) were explicitly excluded, while alanine-alanine (AA) and alanine-valine (AV) combinations were permitted. This strategy yielded 27 discrete variants per group (54 in total), which were generated using the PyMOL Mutagenesis Wizard. The resulting models were subjected to preliminary screening via Rosetta energy minimization using a fixed random seed (11105) to ensure reproducibility of both energy scores and structural conformations.

The top five sequence candidates from each group (ranked by lowest Rosetta energy scores) were further evaluated using the Rosetta FastRelax protocol. Each design was processed across 60 independent replicates (comprising three independent runs of 20 replicates each using random seeds). The resulting distribution of these 60 raw data points for each candidate is shown in **Fig. S3**, demonstrating strong convergence and substantial overlap among the top designs. To assess statistical significance given the small energetic differences between candidates, we performed a bootstrap analysis with 1,000 iterations. In each iteration, 20 samples were drawn with replacement from the pooled 60 replicates to calculate the mean Rosetta Energy Units (REU) and standard deviation.

The final statistical summary of the bootstrap results is provided in **Table S9**. The top five candidates in both groups occupied a highly competitive energetic landscape, with mean scores within 5 REU of one another. Final design selection integrated these quantitative scores with structural intuition. For Group 1, G1_mutant_009 (L6I-L81L-C24A-C98A) was selected (−769.3 ± 0.8 REU) for experimental testing. While this variant does not have the lowest median score, it does have the distinction of having the lowest score observed for any replicate. More importantly, we prioritized the L6I mutation because of the superior β-sheet propensity for isoleucine and because prospective modeling in PyMOL indicated that the 2^nd^ γ-carbon of Ile6 atom effectively refilled the space vacated by the C24A/C98A substitutions, pointing directly into the hydrophobic cavity (**Figs. 3C-3E**). For Group 2, we selected the variant with both the lowest median score and the lowest observed score: G2_mutant_008 (L142L-M175M-C161V-C230A with −770.1 ± 0.7 REU). Beyond yielding the most favorable REU score this variant maintained the heavy atom count, added a β-branched amino acid with β-sheet propensity, and demonstrated high quality internal packing (**Figs. 3F-3H**). In conclusion, this conservative protein design workflow produced a disulfide-free scaffold predicted to remain structurally robust while eliminating oxidation-prone residues. All scripts used for this computational pipeline and the associated analysis are available at github.com/Ashlyn303/disulfide-replace.

### Cα Root-Mean-Square Deviation (RMSD_Cα_) analysis

Root-Mean-Square Deviation (RMSD) values were calculated using the super command in PyMOL (v3.1.3, Schrödinger, LLC). To focus on backbone conformational differences, calculations were restricted to the Cα atoms of the VH (EVKLV…LTVSS) and VL (DIVLT…LEIKR) domains. The flexible (GGGGS)_3_ linker and the C-terminal tail (AAAKGEFGGGGGSGGGGSENLYFQ) were excluded to prevent high-mobility regions from biasing global metrics. All reported RMSD_Cα_ values reflect a simultaneous global rigid-body alignment of both domains. To ensure a comprehensive representation of structural deviation, calculations were performed with zero cycles of outlier rejection (cycles=0), ensuring all defined Cα atoms contributed to the final reported values.

### Fluorescence recovery after photobleaching

For FRAP imaging, U-2 OS cells seeded in MatTek chambers were transiently transfected with 1.25 µg of plasmid encoding mEGFP-tagged intrabody, together with 1.25 µg of plasmid encoding the corresponding 1x epitope tag fused to mCh-H2B. FRAP experiments were performed 20 h post-transfection.

FRAP data were analyzed using a custom Python pipeline that automatically reads microscope data files (.lif) via a custom loader to extract image stacks, time intervals, and pixel size metadata. The pipeline first concatenates the multi-part acquisitions, that is, pre-photobleaching images (Pre), post-photobleaching images acquired at a 1 frame/s (fps; Pb1), and post-photobleaching images acquired at a 0.2 fps (Pb2) into a single time-lapse movie. Subsequently, the Cellpose library ^43^ was used to segment the cell nucleus. The pipeline automatically identifies the FRAP ROI by detecting the bleached spot between the pre- and post-photobleaching frames, or by using a threshold-based approach to track its intensity across all frames. Additional intensity measurements were collected for the nucleus, background, and pseudo-cytosolic regions. Fluorescence intensities within the FRAP ROI were background-subtracted, min-max normalized, and reported as time-dependent recovery curves. For datasets in which fluorescence recovered to near pre-bleach levels, recovery kinetics were fitted to a single-exponential model to extract half-times (t_1/2_) and goodness-of-fit (r^2^) values. ROIs with poor detection or tracking were excluded following manual inspection.

### Measuring binding affinity of intrabody using isothermal titration calorimetry

Isothermal titration calorimetry (ITC) was used to quantify the binding affinity of anti-UTag-IB and its cysteine-free variant (anti-UTag-IB(ΔCys)) to UTag (GGSGGMSLPGRWKPKMGGSGG, GenScript). Experiments were conducted at 25 °C using a MicroCal ITC200 calorimeter (Malvern Panalytical). Prior to ITC measurements, purified intrabody proteins were dialyzed overnight against PBS (pH 7.4) and diluted to 10 µM in the dialysis buffer. The UTag peptide was dissolved in the same dialysis buffer at 100 µM to minimize heats of dilution. The sample cell (∼300 µL) was loaded with the intrabody and the syringe was filled with the UTag peptide. Each titration consisted of an initial 0.4 µL injection followed by 18 successive 2.0 µL injections, with a 180 s spacing between injections to allow the signal to return to baseline. The stirring speed was set to 750 rpm. Reference titrations of UTag peptide into the dialysis buffer were performed under identical conditions to correct for heats of dilution.

Raw thermograms were integrated, corrected for the heat of dilution, and analyzed using NITPIC software^44^. The corrected heats were plotted as a function of the peptide-to-protein molar ratio and fitted to a one-site binding model using nonlinear least-squares regression. Three independent titrations were performed for each intrabody variant, and each replicate was fitted individually to obtain the stoichiometry (n), binding enthalpy (ΔH), and association constant (K_a_). The dissociation constant was calculated as K_D_ = 1/K_a_. Reported K_D_ values represent the mean ± SEM from three independent experiments.

### Thermostability characterization of anti-UTag intrabodies

Rosetta (DE3) competent *E. coli* cells (Sigma-Aldrich) were transformed with pET23b plasmid encoding wild-type anti-UTag-scFv, anti-UTag-IB, or anti-UTag-IB(ΔCys), each fused to a TEV cleavage site and a C-terminal mEGFP. Single colonies were grown at 37 °C and 225 rpm until OD600 reached 0.5. Protein expression was induced by adding 0.5 mM IPTG (RPI), followed by incubation at 16 °C and 225 rpm for 20 h. Cells were harvested by centrifugation and lysed by sonication at 4 °C (60% amplitude, 2 s on/30 s off cycles, 60 cycles total; Diagenode) in lysis buffer containing 50 mM Tris, 150 mM NaCl, 10 mM imidazole, 5% glycerol, and Complete protease inhibitor (Roche). Insoluble debris was removed by centrifugation at 12,000 x g for 45 min at 4 °C, and the clarified lysate was transferred to a fresh tube. Equal volumes of clarified lysate were incubated for 15 min at 4 °C, 50 °C, 60 °C, or 70 °C, followed by centrifugation at 20,000 x g for 15 min at 4 °C. The supernatant was collected, clarified by a second centrifugation step, and subsequently analyzed by western blots.

### Determining the melting temperature of intrabodies using circular dichroism

Purified anti-UTag-IB and anti-UTag-IB(ΔCys) proteins were diluted to 10 µM in PBS (pH 7.4) and subjected to circular dichroism (CD) analysis. Thermal denaturation was monitored by recording ellipticity at 216 nm while increasing the temperature from 5 to 95 °C. Each experiment was performed in triplicate for both proteins. The resulting data were fitted to a sigmoidal model using scipy.optimize.curve_fit in Python to determine the melting temperature (T_m_) defined as the temperature at which 50% of the protein population is unfolded. Reported T_m_ values represent the mean ± SEM from three independent experiments.

### Western blots

For the thermostability characterization, 40 µL of each lysate was mixed with SDS-PAGE sample buffer, incubated at 95°C for 10 min, and loaded to a 4-20% Mini-PROTEAN TGX Gel (Bio-Rad). The SDS-PAGE gel was run for 5 min at 150 V and 40 min at 200 V for proper protein separation. Proteins were transferred onto a PVDF membrane (Bio-Rad) and blocked for 1 h with a blocking buffer (5% non-fat milk powder in TBST (0.05% Tween-20 (Thermo Scientific) in TBS buffer, pH 7.4)). The blot was incubated with mouse monoclonal anti-GFP antibody diluted in the blocking buffer (1:5000; Cell Signaling Technology) overnight at room temperature with gentle agitation. After washing the blot six times with TBST, the blot was probed with horseradish peroxidase (HRP)-conjugated anti-mouse antibody diluted in the blocking buffer (1:5000; Cell Signaling Technology) for 1 h at room temperature with gentle agitation. Following six additional washes with TBST, signals were developed using Clarity™ Western ECL substrate (Bio-Rad) and imaged on a ChemiDoc MP imaging system (Bio-Rad).

For visualization of tagged KDM5B reporters, U-2 OS cells seeded in 35 mm dishes were transiently transfected with the indicated constructs and harvested 12 h post-transfection. Cell pellets were lysed in 150 µL Pierce RIPA buffer (Thermo Scientific) supplemented with Complete protease inhibitor (Roche). Total protein concentration was determined using a Pierce BCA protein assay kit (Thermo Scientific). Equal amounts of protein (50 µg) were mixed with SDS–PAGE sample buffer, incubated at 95 °C for 10 min, and loaded onto a 4–15% Mini-PROTEAN TGX gel (Bio-Rad). Electrophoresis was carried out and proteins were transferred to blots as described above. Blots were processed as described above, except that blots were incubated with rabbit polyclonal anti-JARID1B antibody (1:1000; Cell Signaling Technology) and rabbit polyclonal anti-tubulin antibody (1:5000; Cell Signaling Technology) as primary antibodies, followed by HRP-conjugated anti-rabbit secondary antibody (1:5000; Cell Signaling Technology). Signals were detected using Western Lightning™ Plus chemiluminescence reagent (Revvity) and imaged on a ChemiDoc MP imaging system (Bio-Rad).

Protein band intensities from Western blots were quantified using the area-under-the-curve method in ImageJ (v1.54p). Raw .TIFF images were imported, and individual lanes were defined using the rectangular selection tool and analyzed via the *Gels* function under the *Analyze* menu. After selecting all lanes, intensity profiles were generated for each lane. Peaks corresponding to bands of interest were identified, and the area under each peak was quantified using the wand tool. For thermostability analysis, band intensities were normalized to the corresponding control samples maintained at 4 °C.

### KDM5B localization imaging

For visualizing 24xUTag and 24xALFA tagged KDM5B localization, U-2 OS cells seeded in MatTek chambers were transiently transfected with each tagged KDM5B plasmid (17 µL of 100 ng/µL) and its corresponding mEGFP fused intrabody (8 µL of 100 ng/µL). For 24xSunTag tagged KDM5B, we transfected 20 µL of the KDM5B plasmid (100 ng/µL) and 5 µL of the anti-SunTag-IB-mEGFP (100 ng/µL). Cells were imaged 24 h post-transfection.

### Translation tracking

To image single-mRNA translation, U-2 OS cells were seeded into MatTek chambers at approximately either 25% confluency 48 h before imaging or 50% confluency 24 h before imaging. For 24xUTag-KDM5B-24xMS2 and 24xALFA-KDM5B-24xMS2 reporters, cells were transiently transfected with reporter plasmid (7.5 µL of 100 ng/µL), the corresponding mEGFP fused intrabody plasmid (2.5 µL of 100 ng/µL), and tdMCP-CAAX plasmid (2.5 µL of 100 ng/µL) were transiently transfected into U-2 OS cells. Imaging was performed approximately 20 h post-transfection. For 24xSunTag-KDM5B-24xMS2 reporter, cells were transiently transfected with the reporter plasmid (15 µL of 100 ng/µL), anti-SunTag-IB-mEGFP plasmid (5 µL of 100 ng/µL), and tdMCP-CAAX plasmid (5 µL of 100 ng/µL) and imaged approximately 4 h post-transfection.

For autocorrelation analysis, the 3D time-lapse movies were acquired at 5 s interval for 360 frames. For harringtonine run-off assays, 4-5 cells per experiment were pre-selected and imaged using a multi-position microscope setup at 1 min per frame. Harringtonine (Cayman; 3 µg/mL) was added after the fifth frame, and imaging continued for an additional 25 min.

### Basic image processing for translation tracking

Microscopy data consisted of five-dimensional image stacks encompassing three spatial dimensions (x, y, z), a fluorescence channel dimension (translation spot signal), and a temporal dimension (t). Typical acquisition parameters included 9 z-slices, 130 nm xy pixel size, 300 nm z-step size, and 16-bit intensity resolution. Multiple time-lapse movies were collected per experimental day. Initial quality control was performed by manually selecting movies in which cells remained viable throughout imaging, stayed in focus, lacked aggregation artifacts, and exhibited detectable translation spots.

Cell segmentation was performed using either a watershed-based algorithm or a deep learning–based approach implemented in Cellpose^43^. In cases where automated segmentation failed, the cell boundary was manually delineated. Final segmentation masks were generated by inspecting multiple frames and expanding the region to encompass the maximal observed cell area, ensuring consistent segmentation across all time points.

Photobleaching correction was performed by calculating the median fluorescence intensity across the entire image of each frame and fitting an exponential decay function to the resulting time series to determine the bleaching rate. Each frame was then normalized by the value of the fitted decay function at time *t*. To avoid introducing artifacts, time-lapse movies exhibiting less than a 5% decrease in fluorescence intensity relative to the first frame were not subjected to photobleaching correction.

### Particle detection and tracking

Particle detection, tracking, and fluorescence intensity quantification were performed using MicroLive^45^. For each time frame, a maximum intensity projection across all z-planes was generated. Diffraction-limited particles were detected and linked into trajectories using an expected spot size of 5 pixels, a search radius of 10 pixels, and a memory of 1 frame to reconnect particles that transiently disappeared.

Spot fluorescence intensities were quantified using MicroLive’s local background subtraction method, in which the mean intensity of a surrounding annular region was subtracted from the mean intensity of the central spot.^45^ Apparent spot sizes were determined by fitting a two-dimensional Gaussian function to each detected spot, optimizing the peak amplitude and spatial standard deviations in the x and y dimensions. Spot size was then estimated as the full width at half maximum (FWHM) calculated from the mean of these standard deviations; failed fits were recorded as not-a-number (NaN) values. Finally, a lower-bound estimate of the signal-to-noise ratio (SNR) was computed for each spot as the ratio of background-subtracted mean intensity to the standard deviation of the local background.

### Harringtonine run-off assay analysis

Images were collected for an initial five frames at a frame rate of 1 frame per min. Subsequently, Harringtonine was applied, and acquisitions were collected for an additional 25 frames post-treatment, making a total of 30 frames with a frame rate of 1 frame per min. Image processing was implemented using a custom Python pipeline. Briefly, as the Harringtonine application involves adding the drug volume to the cell media during the microscopy session, this may move the field of view out of focus; to address this, image processing was performed using maximum projection across all z-planes. Whole-cell segmentation was implemented using the previously described methods. Given the slow framing rate and the small number of frames collected, we did not observe a decrease in the full-image intensity, and photobleaching correction was unnecessary. Translation spots were detected with TrackPy using a user-defined threshold, and spot detection was implemented for each frame in the video.^46^ As the number of detected spots decreased over time after the Harringtonine application, we used the average number of spots detected before the Harringtonine application to calculate the average intensity for the rest of the movie. This correction was necessary to account for the spots disappearing from the measurements. The final intensity vector representing the mean of the detected spots in a single cell is given by:

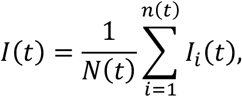

where

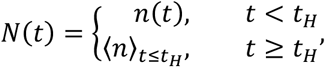

and ⟨*n*⟩ denotes the temporal average of *n*(*t*) the time prior to the harringtonine application (*t*_*H*_).

To represent the intensity values on comparable scales, the final intensity values in run-off plots were normalized a second time using the frames before treatment. This step was necessary to account for imaging and basal intensity heterogeneity.

### Autocorrelation Function Analysis

Data were collected at a frame rate of 5 s for a total of 360 frames. 2D particle tracking was implemented as described before. Trajectories comprising fewer than 30% of the total frames were excluded from further analyses to eliminate short, potentially spurious tracks. As described by Coulon et al.^47^, signals with a low signal-to-noise ratio (SNR<0.5) were discarded. Fluorescence Correlation Spectroscopy (FCS) was performed according to established protocols^39,48^. The autocorrelation functions were calculated as follows:

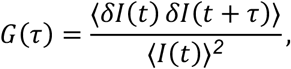

where *I*(*t*) is the fluorescence intensity at time *t*, ⟨⋅⟩ denotes the temporal average, *δI*(*t*) is defined as follows *δI*(*t*) = *I*(*t*) − ⟨*I*(*t*)⟩, and *τ* represents a discrete time interval shift.

Following Coulon *et al*.^39^, a bootstrapping approach was employed to estimate the uncertainty in the autocorrelation function, additionally, the autocorrelation at zero lag (*G*(*0*)) was replaced with a linear extrapolation from the 5 adjacent time points to minimize shot noise artifacts. Finally, the reported mean autocorrelation function values were shifted to zero to account for basal values.

To calculate the final average correlation, outlier trajectories were removed to avoid bias from extreme values resulting from particle-tracking or intensity-quantification errors. Trajectories with mean correlations deviating more than four times the median absolute deviation (MAD) from the overall median were classified as outliers and excluded, removing approximately 15-20% of the trajectories.

To test the accuracy of our correlation calculation method, we used the simulated intensities generated with the mathematical model described in the next section and confirmed that we could effectively recover the elongation and initiation rates used in the simulation after applying the same pre-processing as in the experimental data (**Figs. S8C-D**).

### Modeling Approach

Modeling is based on the TASEP (totally asymmetric exclusion process)^37,49^. The model was solved stochastically using Monte Carlo-like simulations^50^. In short, the model describes ribosome initiation, elongation, and termination processes. Elongation rates are adjusted to reflect codon usage in the human genome (*k*_*e*_(*i*) = *k̅*_*e*_(*u*(*i*)/*u̅*), *i* indicates a specific codon), 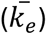 is the average elongation constant, *u*(*i*) represents the frequency of codon *i*, *u̅* is the average frequency of all amino acid encoding codons, and a ribosome footprint (*nf*) of 9 codons guarantees that initiation and elongation only occur if the codons downstream are unoccupied.

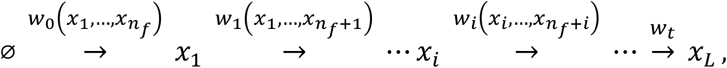

Where 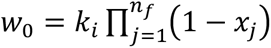 with initiation rate 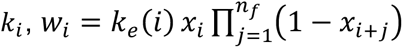, and the termination rate is assumed to be equal to the average elongation constant, 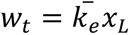.

The TASEP model simulates temporal ribosome occupancy. From this, a simulated fluorescence intensity vector *I*_*sim*_ was constructed using a vector containing the position of the tag in the original DNA sequence, as follows:

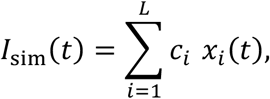

where, *c*_*i*_ indicates the number of epitopes (bound by fluorescent probes) that accumulate by the time translation reaches the codon *i*, and *L* is the gene length. This modeling strategy has been described in detail by Aguilera et al. ^37^

### Data fitting and parameter estimation

We calculate the likelihood of observing the experimental data *D* given a parameter set *θ*. For this, we assume that at each lag time *τ*, the experimental correlation function *G*_*exp*_(*τ*) is normally distributed around the simulated correlation function *G*_*sim*_(*τ*, *θ*), with variance estimated from the experimental uncertainty at the lag time. As an approximation, we treated uncertainty at each lag independently. In other words:

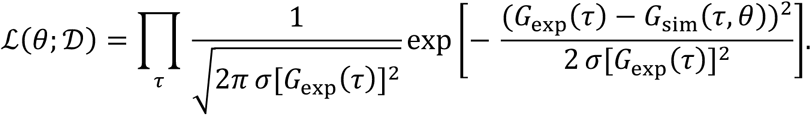

Taking the logarithm, the log-likelihood becomes

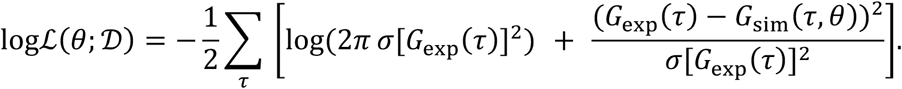

For the optimization process, we simplified the objective function by combining independent terms of parameters *θ* and scaling factors. Thus, the negative log-likelihood can be approximated by a nonlinear least-squares fit, as described by Coulon *et al.*^39^:

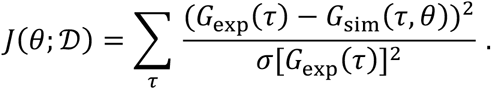

Optimization methods were implemented to obtain the parameter set that minimizes the proposed objective function, that is:

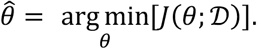

Because our translation model is nonlinear, the optimization problem cannot be solved analytically. Instead, we performed an exhaustive grid search over the initiation and elongation rates. Specifically, we sampled

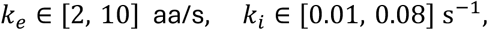

at 80 evenly spaced values each, yielding 6,400 total parameter combinations. The objective function was evaluated at every grid point to identify the optimum (**Figs. S9A-S9D**). These bounds were chosen to reflect biologically plausible rates as established previously ^37^.

To obtain robust point estimates, we report the *median* of the top 5% lowest-cost grid points. Gaussian smoothing was applied to the cost surface to reduce noise. To characterize the sensitivity of the cost surface near the optimum, we identified and calculated the 90% confidence interval, defined by the 5th and 95th percentiles of the parameter distributions within the top 5% of this population.

To extract elongation rates from the harringtonine data, we optimized TASEP model parameters by minimizing the difference between model predictions and experimental data. For this, we ran TASEP simulations in which ribosomal initiation was reduced by 95% after 5 minutes, matching the experimental time of harringtonine application. To account for the diffusion time of the drug to the cell, a 30-second delay was incorporated before the onset of inhibition. Elongation rates were then sampled over the range of 1.0-8.0 aa/s using 15 evenly spaced test values, and the initiation rate was fixed at 0.03 s⁻¹. For each candidate elongation rate, model-data agreement was quantified using chi-squared over the harringtonine response window (t = 0-24 min).

### Quantifying Translation Kinetics from the Model

Initiation and elongation were directly estimated by fitting the TASEP model to the ACF analysis and harringtonine run-off assays as described in the previous section. Additionally, we estimated the average number of ribosomes (*R*, defined as the number of ribosomes per mRNA), and the ribosomal density (*ρ*, defined as the fraction of the mRNA decoding codons occupied by ribosomes), following Lamberti *et al.*, definitions^38^. For *R*, we used:

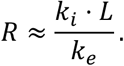

Ribosomal density (*ρ*), defined as the fraction of RNA covered by ribosomes, was approximated and expressed as a percentage as:

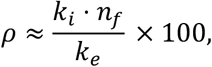

where *n*_*f*_ is the ribosome footprint (reported ∼ 9 codons^51^).

Ribosomal distance (*r*_*d*_), defined as the approximated distance (aa) between consecutive ribosomes assuming homogeneous distributions, was calculated as follows:

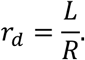

### Statistical analysis

Statistical significance between two groups was analyzed using Mann-Whitney U test from scipy.stats library. Statistical significance was denoted as follows: **** p < 0.0001, *** p < 0.001, ** p < 0.01, and * p < 0.05. Only significant differences are indicated in the figures. The numbers of cells, trajectories, and independent experimental replicates are reported in the figures and figure legends.

## Competing Interests

The authors declare no competing interests.

## Acknowledgments

We thank Robb Welty at the University of Colorado Anschutz Biophysics Shared Resource Facility for collecting and analyzing the ITC and CD data. We thank Hataichanok (Mam) Scherman at the Colorado State University Protein Expression and Purification Facility for purifying the anti-UTag intrabodies. This work was supported by NIH grants K99GM141453 (N.Z.), R00GM141453 (N.Z.), R35GM160021(N.Z.), 1R01AI168459-01A1 (C.S.) and Cystic Fibrosis Foundation grant 005749A123 (N.Z.).

## Data Availability

All data and microscopy movies will be made publicly available upon publication. Crystal structures of anti-UTag intrabodies have been deposited in the Protein Data Bank under accession codes 7URL, 9N97, and 8V8F.

## Code Availability

Source code for cysteine-free intrabody design is available at https://github.com/Ashlyn303/disulfide-replace. Source code for computational modeling, TASEP simulations, parameter optimization, and microscopy data analysis is available at https://github.com/ningzhaoAnschutz/utag_paper. Image processing was performed using MicroLive, for which the source code is available at https://github.com/ningzhaoAnschutz/microlive.

## Author contributions

N.Z., T.S., C.S., and B.G. conceived the experiments. N.Z., L.A., S.C., J.Y., R.S., and H.O. conducted the experiments. S.C., J.D., and C.S. designed the cysteine-free intrabody; N.Z., L.A., S.C., C.S., and R.S. analyzed the data; L.A., S.C., R.S., C.S., and N.Z. drafted the manuscript. All authors reviewed the manuscript.

## Notes

### Competing Interest Statement

The authors have declared no competing interest.

