## Supplemental File for "UTag, a cysteine-free thermostable tagging system for tracking single mRNA translation live"

Supplementary Figures S1-S9

Supplementary Tables S1-S9

Supplementary Movies S1-S4

**Supplementary Movie S1: Translation tracking using the UTag tagging system.** A representative cell expressing anti-UTag-IB-mEGFP and 24xUTag-KDM5B-24xMS2 to visualize translation. The movie is related to Figs. 7C-7E. Field of view is 66.51 x 66.51 µm. 5 s interval.

**Supplementary Movie S2: Translation tracking using the cysteine-free UTag tagging system.** A representative cell expressing anti-UTag-IB(ΔCys)-mEGFP and 24xUTag-KDM5B-24xMS2 to visualize translation. The movie is related to Figs. 7C-7E. Field of view is 66.51 x 66.51 µm. 5 s interval.

**Supplementary Movie S3: Translation tracking using the SunTag system.** A representative cell expressing anti-SunTag-IB-mEGFP and 24xSunTag-KDM5B-24xMS2 to visualize translation. The movie is related to Figs. 7C-7E. Field of view is 57.67 x 50.53 µm. 5 s interval.

**Supplementary Movie S4: Translation tracking using the ALFA-tag tagging system.** A representative cell expressing anti-ALFA-IB-mEGFP and 24xALFA-KDM5B-24xMS2 to visualize translation. The movie is related to Figs. 7C-7E. Field of view is 66.51 x 66.51 µm. 5 s interval.

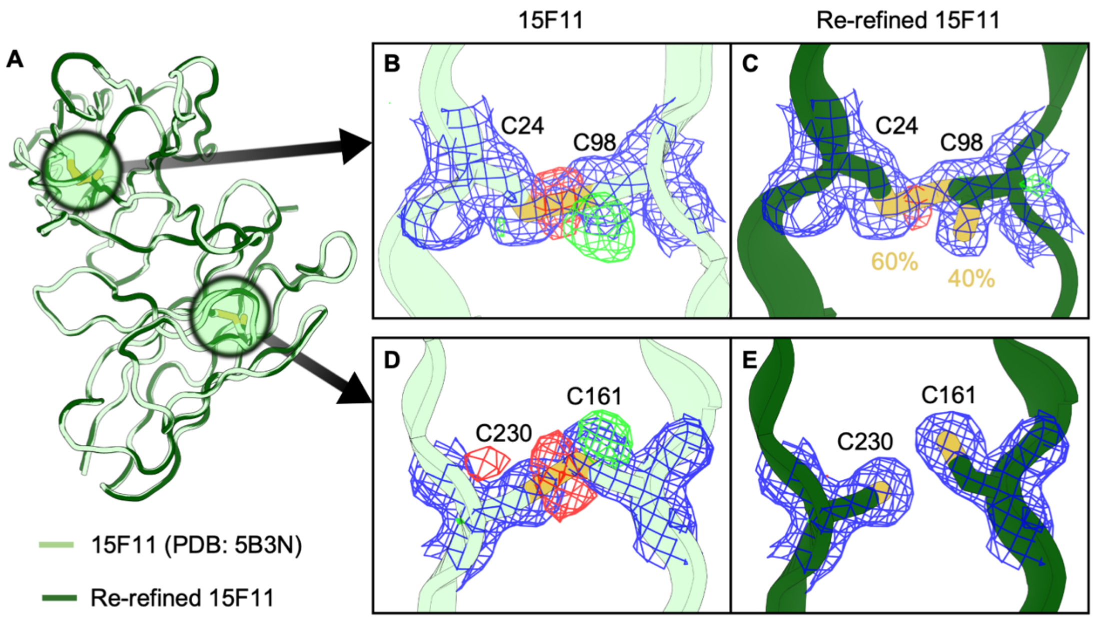


**Figure S1. Structural comparison of 15F11 scFv (PDB: 5B3N) before and after re-refinement. (A)** Structural superposition of the original RCSB PDB: 5B3N entry (light green) and the re-refined model (dark green). **(B–C)** Electron density and difference Fourier maps centered on the first disulfide bond (C24-C98) for the original (B) and re-refined (C) structures. **(D–E)** Close-up views of the second potential disulfide bond for the original (D) and re-refined (E) models. Electron density is shown as a $2F_O$-$F_C$ map (blue mesh, contoured at 1.0 σ), while model discrepancies are highlighted by the $F_O$-$F_C$ difference map (contoured at ± 3.0 σ). Green density indicates positive $F_O$-$F_C$ peaks where additional atomic model may be warranted. Red density indicates negative $F_O$-$F_C$ peaks where the model over-represents the experimental density.

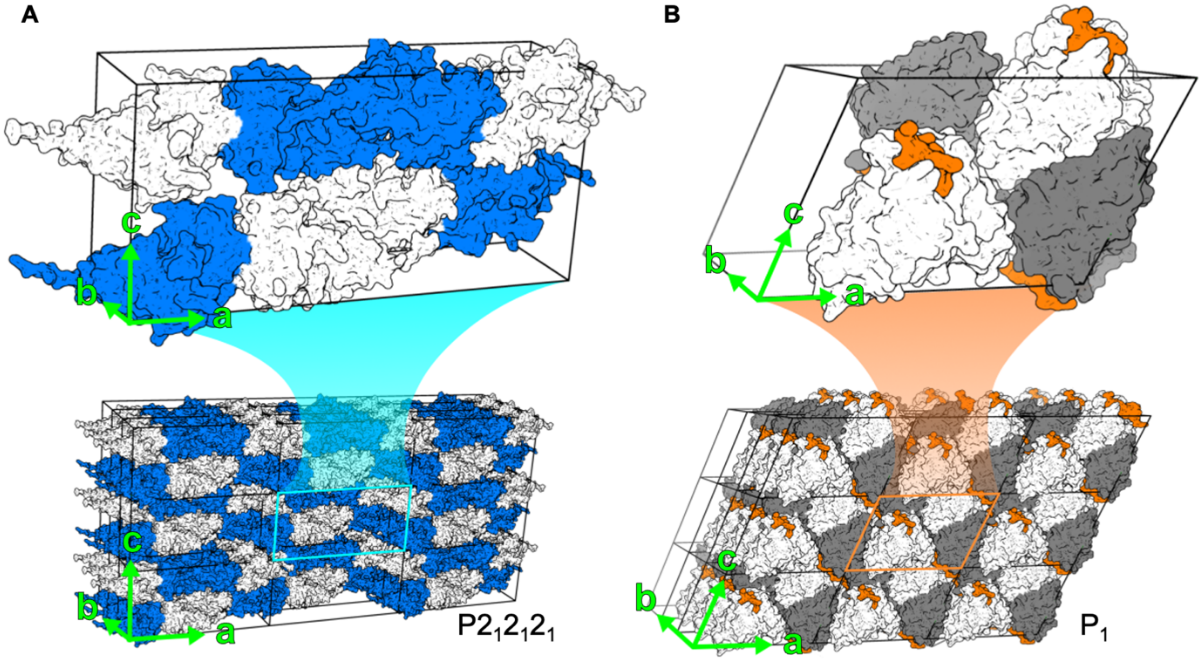


**Figure S2. Supercell structures of anti-UTag intrabody with and without UTag. (A)** Anti-UTag-IB (PDB: 7URL) arranged in a 3 x 3 x 3 unit-cell supercell containing 216 protein chains. Each unit cell contains four copies of two asymmetric molecules with space group $P2_12_12_1$. The two asymmetric copies of protein chains are shown in blue and white. **(B)** Anti-UTag-IB/UTag complex (PDB: 8V8F) arranged in a 3 x 3 x 3 unit-cell supercell containing 108 protein chains and 108 peptide chains. Each $P_1$ unit cell contains four copies of intrabody-peptide complexes. Protein chains are shown in gray and white. UTag peptide is in orange. Both panels show the crystal packing with crystallographic axes labeled (a, b, c in green).

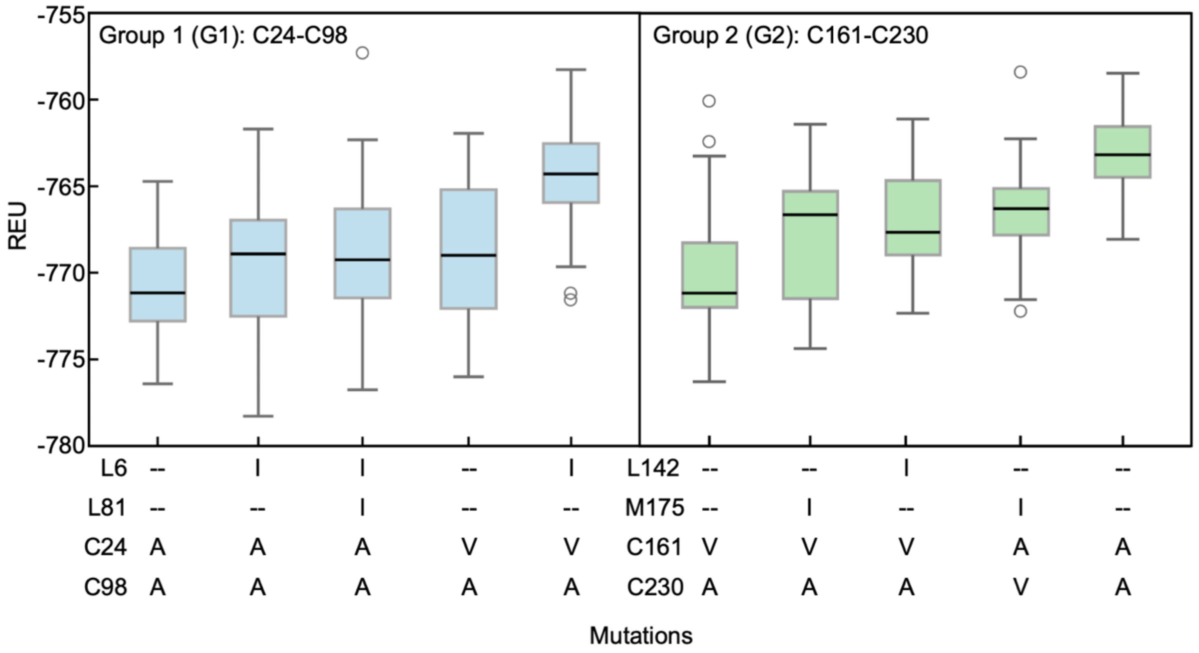


**Figure S3. Computational stability analysis of cysteine-free intrabody variants.** Rosetta Energy Units (REU) for the top five candidates in **(A)** Group 1 (C24–C98) and **(B)** Group 2 (C161–C230). The cysteine residues are numbered according to the sequence of 15F11 (PDB: 5B3N). Data represent 60 independent FastRelax replicates per design. Boxes indicate the interquartile range (IQR), and the central line represents the median. The maximum and minimum values observed within 1.5 x IQR are shown as whiskers (error bars), with circles indicating statistical outliers defined as points falling outside this range.

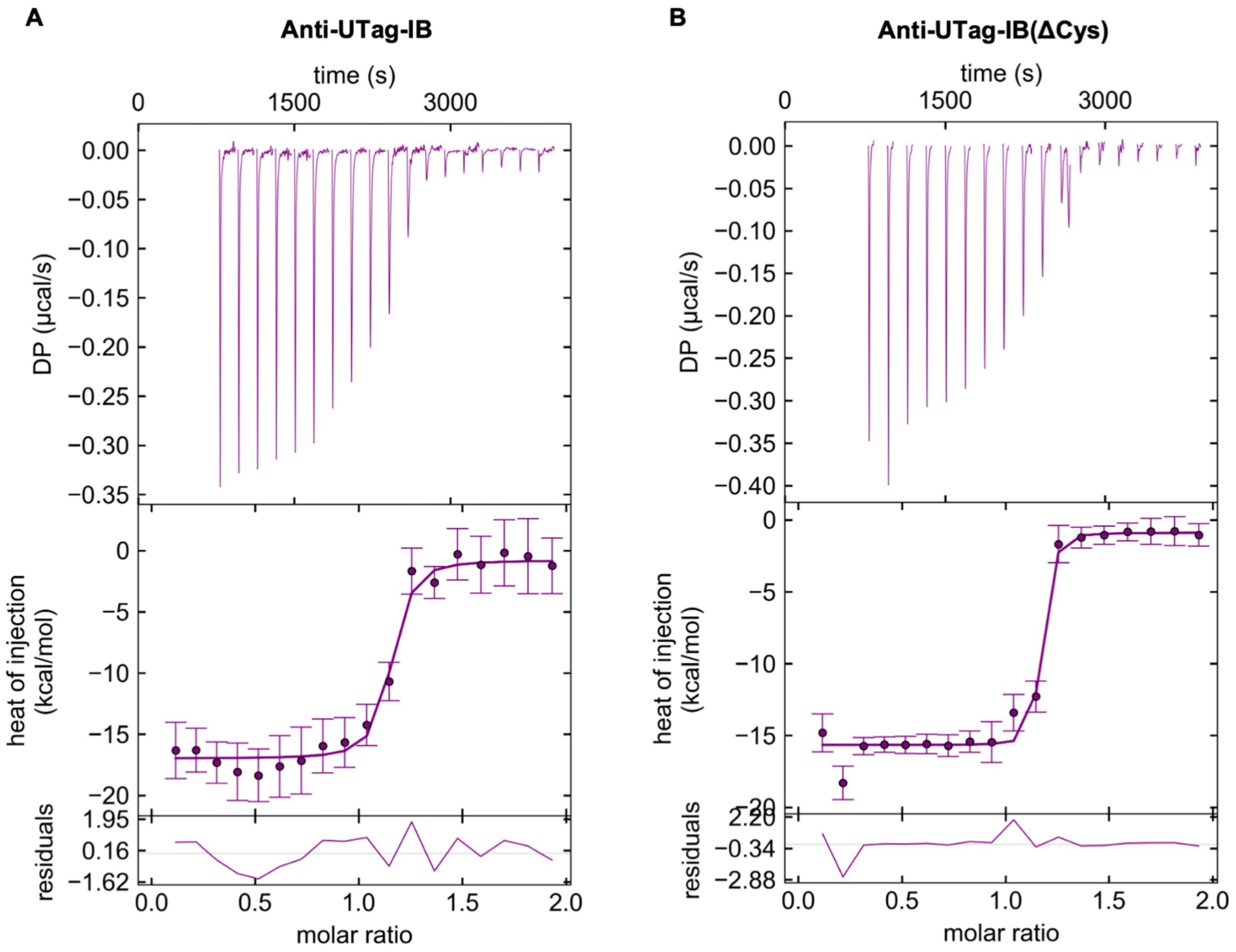


**Figure S4. Anti-UTag-IB and anti-UTag-IB(ΔCys) bind to UTag with low nM binding affinity. (A)** Representative isothermal titration calorimetry (ITC) experiment of anti-UTag-IB. **(B)** Representative ITC experiment of anti-UTag-IB(ΔCys). For each intrabody, the top panel shows the raw thermogram, the middle panel shows the integrated injection heats (kcal/mol) versus molar ratio fitted to a single-site sigmoid binding model (solid line), and the bottom panel shows the residuals of the fit. For each intrabody, the ITC experiment was repeated three times.

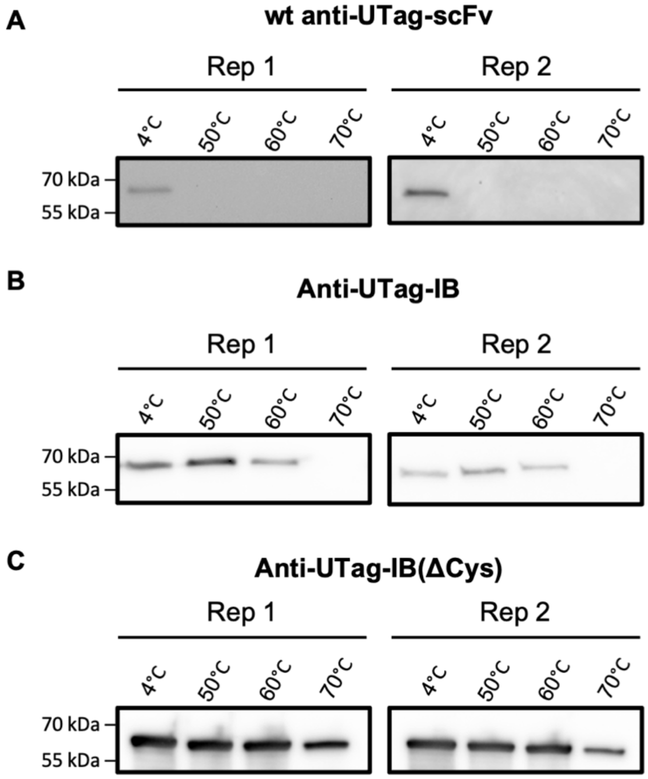


**Figure S5. Western blots of characterizing the thermostability of anti-UTag intrabodies.** Two independent western blots of soluble fractions after incubation for 15 min at 4 °C, 50 °C, 60 °C, and 70 °C for **(A)** wild-type anti-UTag-scFv, **(B)** anti-UTag-IB, and **(C)** anti-UTag-IB(ΔCys).

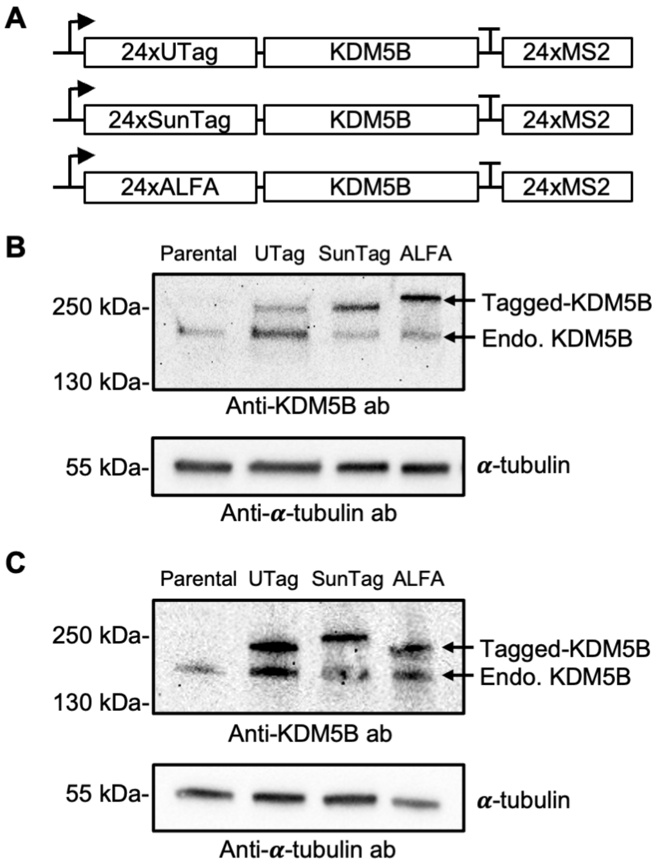


**Figure S6. Validation of full-length KDM5B expression with different tagging systems. (A)** Schematic illustration of reporter constructs containing 24xUTag-, 24xSunTag-, or 24xALFA-tagged KDM5B. **(B)** Western blot analysis of the reporter constructs shown in (A). **(C)** Independent replicate of the western blot analysis shown in (B).

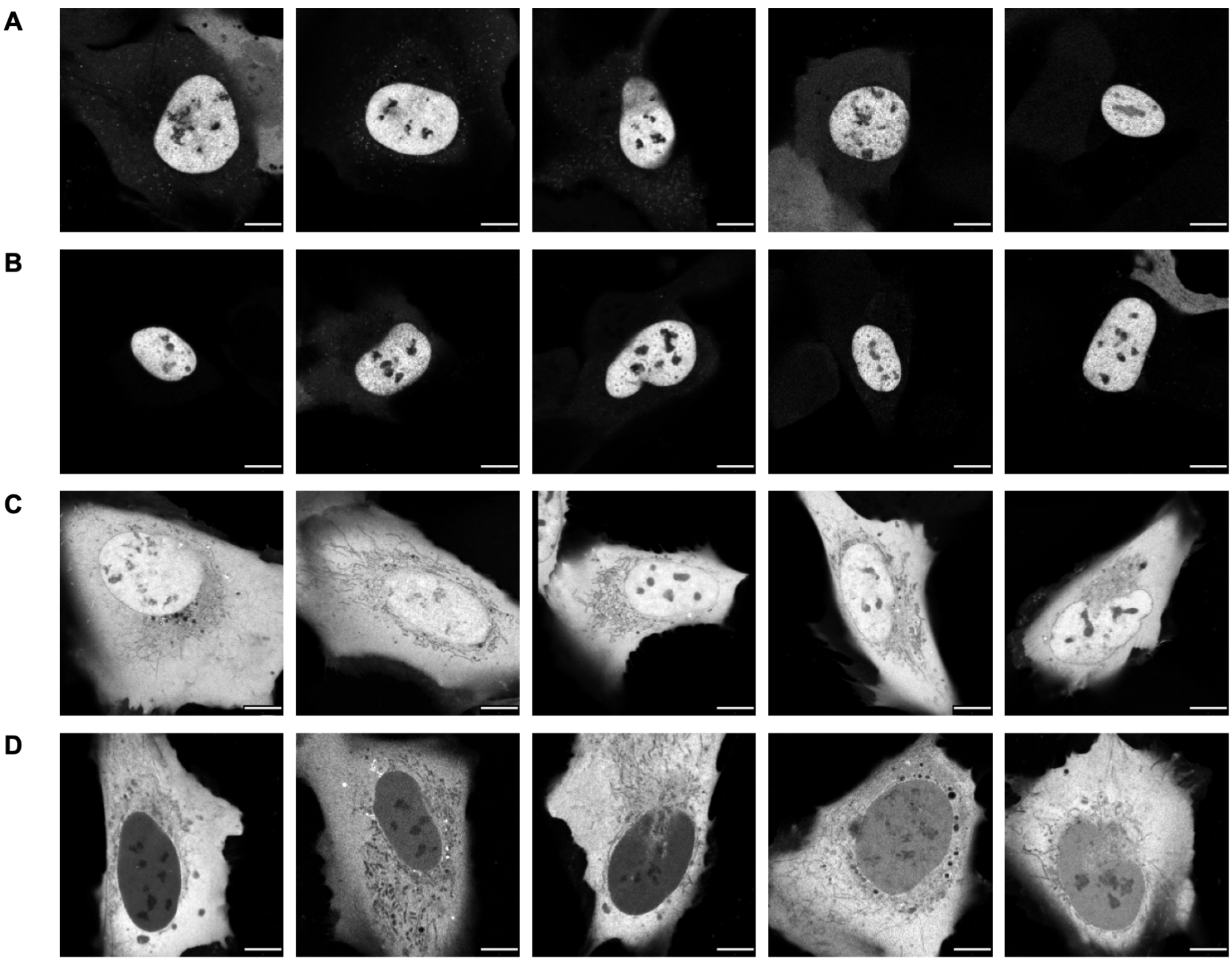


**Figure S7. Cellular localization of tagged-KDM5B visualized by cognate fluorescent intrabodies.** Representative cell images showing different tagged KDM5B cellular localization 24 h post-transfection. **(A)** Anti-UTag-IB. (B) Anti-UTag-IB(ΔCys). (C) Anti-SunTag-IB. (D) Anti-ALFA-IB. Scale bars: 10 µm.

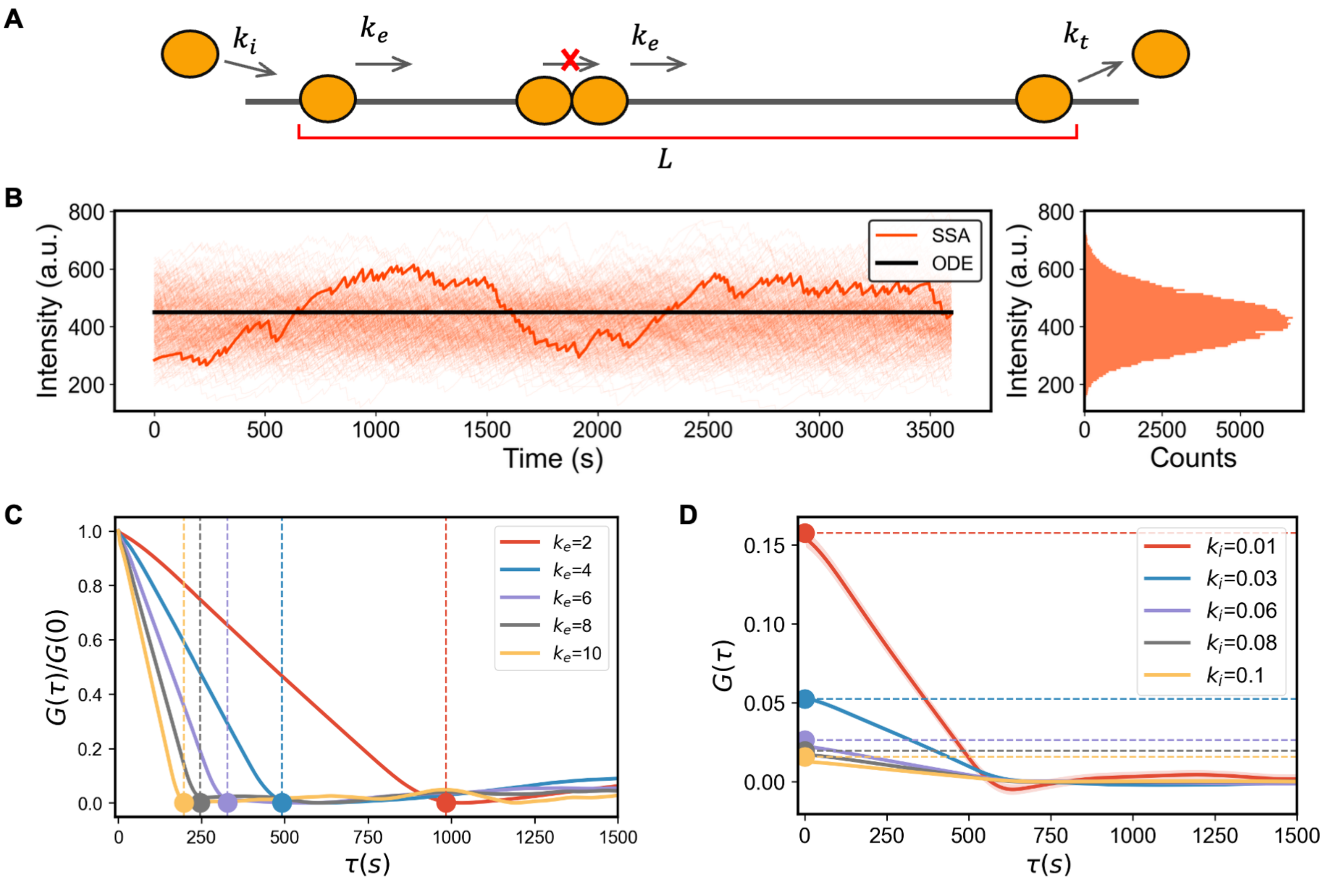


**Figure S8. TASEP modeling simulates single-mRNA translation. (A)** Schematic illustrating key steps and main factors affecting translation included in the TASEP model, including initiation ($k_i$), elongation ($k_e$), gene length ($L$), ribosome exclusion ($n_f$), and termination ($k_t$). **(B)** The TASEP model was solved using 300 repetitions in the Gillespie Algorithm, using representative parameters obtained from the parameter optimization process ($k_e$ = 3.1 aa/s and $k_i$ = 0.031 s$^{-1}$). **(C)** Normalized autocorrelation function (ACF) calculation from stochastic simulations using a parameter sweep for $k_e \in [2, 10]$ aa/s (solid lines). **(D)** ACF calculation from stochastic simulations using a parameter sweep for $k_i \in [0.01, 0.1]$ s$^{-1}$ (solid lines). (C, D) Dashed lines show the corresponding approximated analytical solutions derived from the closed-form expressions for $k_i = 1/\,(G(0)\,\tau)$ and $k_e = L/\tau$ , where $\tau$ represents the dwell time.

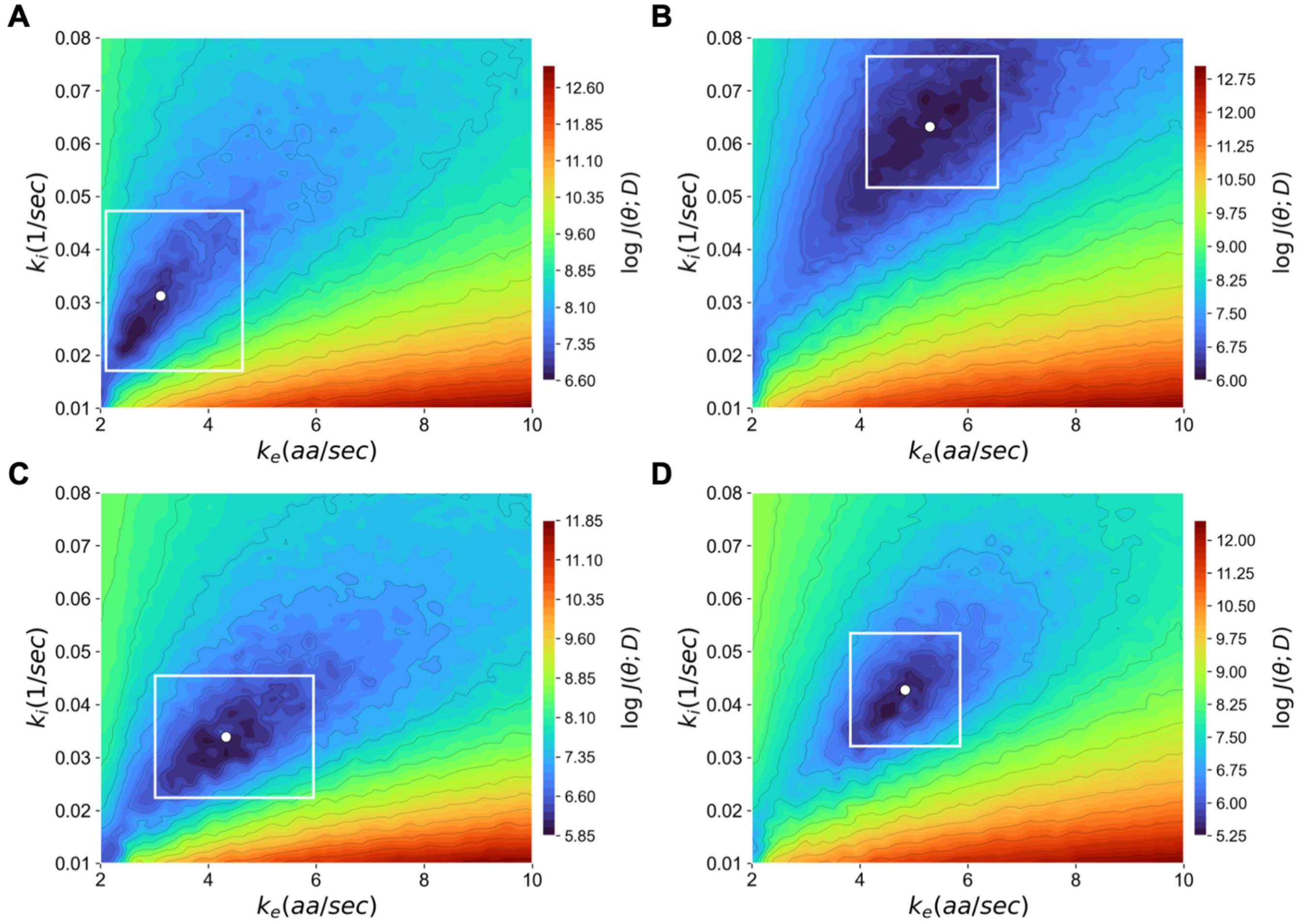


**Figure S9. Parameter optimization strategy using an 80 x 80 grid.** For a better visualization, the color map is plotted using the log of the objective function. The parameter estimates (white dot) represent the median k$i$ and k$e$ values from the top 5% of grid search solutions ranked by cost. The rectangle delineates the 90% confidence interval, defined by the 5th and 95th percentiles of the parameter distributions within the top 5% of this population. **(A)** Anti-UTag-IB-mEGFP. **(B)** Anti-UTag-IB(ΔCys)-mEGFP. **(C)** Anti-SunTag-IB-mEGFP. **(D)** Anti-ALFA-IB-mEGFP.

**Table S1. Structure wwPDB validation metrics comparison of PDB 5B3N vs re-refined 5B3N.**

| Metric | PDB 5B3N | Re-Refined |
|---|---|---|
| $R_{free}$ | 0.253 | 0.234 |
| Clash score | 3 | 1 |
| Ramachandran outliers | 0 | 0 |
| Sidechain outliers | 2.1% | 0 |
| RSRZ outliers | 3.0% | 1.3% |

**Table S2. $RMSD_{C\alpha}$ comparing the local structural environments of mutated residues between parental and cysteine-free intrabody structures.**

| Mutation | Before | After | $RMSD_{C\alpha}$ (Å) |
|---|---|---|---|
| L4I | VK**L**VE | VK**I**VE | 0.05 |
| C22A | LS**C**AA | LS**A**AA | 0.08 |
| C96A | YY**C**AR | YY**A**AR | 0.13 |
| C163V | IS**C**RA | IS**V**RA | 0.13 |
| C232A | YY**C**QQ | YY**A**QQ | 0.14 |

**Table S3. Phenix.Table_one for anti-UTag intrabody (PDB: 7URL).**
Statistics for the highest-resolution shell are shown in parentheses.

| | |
|---|---|
| **Wavelength** | 1.1 |
| **Resolution range** | 37.03 - 1.411 (1.462 - 1.411) |
| **Space group** | P 21 21 21 |
| **Unit cell** | 41.725 76.12 160.628 90 90 90 |
| **Total reflections** | 176609 (2042) |
| **Unique reflections** | 88732 (1114) |
| **Multiplicity** | 2.0 (1.8) |
| **Completeness (%)** | 89.33 (11.37) |
| **Mean I/sigma(I)** | 8.56 (0.91) |
| **Wilson B-factor** | 16.78 |
| **R-merge** | 0.05586 (0.5636) |
| **R-meas** | 0.079 (0.7971) |
| **R-pim** | 0.05586 (0.5636) |
| **CC1/2** | 0.998 (0.369) |
| **CC*** | 0.999 (0.734) |
| **Reflections used in refinement** | 88713 (1114) |
| **Reflections used for R-free** | 2000 (25) |
| **R-work** | 0.1836 (0.3413) |
| **R-free** | 0.2225 (0.2710) |
| **CC(work)** | 0.974 (0.577) |
| **CC(free)** | 0.967 (0.638) |
| **Number of non-hydrogen atoms** | 6077 |
| **macromolecules** | 5661 |
| **ligands / solvent** | 24 / 392 |
| **Protein residues** | 488 |
| **RMS(bonds) / RMS(angles)** | 0.018 / 1.75 |
| **Ramachandran favored (%)** | 96.65 |
| **Ramachandran allowed (%)** | 3.09 |
| **Ramachandran outliers (%)** | 0.26 |
| **Rotamer outliers (%)** | 5.05 |
| **Clashscore** | 5.38 |
| **Average B-factor** | 27.92 |
| **macromolecules** | 27.16 |
| **ligands** | 70.79 |
| **solvent** | 36.20 |
| **Number of TLS groups** | 14 |

**Table S4. Phenix.Table_one for cysteine-free anti-UTag intrabody (PDB: 9N97).**
Statistics for the highest-resolution shell are shown in parentheses.

| | |
|---|---|
| **Wavelength** | 0.99999 |
| **Resolution range** | 42.9 - 1.45 (1.49 - 1.45) |
| **Space group** | P 21 21 21 |
| **Unit cell** | 40.679 74.521 157.394 90 90 90 |
| **Total reflections** | 167334 (8893) |
| **Unique reflections** | 84215 (4729) |
| **Multiplicity** | 2.0 (1.9) |
| **Completeness (%)** | 98.05 (78.06) |
| **Mean I/sigma(I)** | 11.24 (0.78) |
| **Wilson B-factor** | 16.57 |
| **R-merge** | 0.03664 (0.8496) |
| **R-meas** | 0.05182 (1.201) |
| **R-pim** | 0.03664 (0.8496) |
| **CC1/2** | 0.999 (0.394) |
| **CC*** | 1 (0.752) |
| **Reflections used in refinement** | 84182 (4718) |
| **Reflections used for R-free** | 2000 (113) |
| **R-work** | 0.1712 (0.3428) |
| **R-free** | 0.2120 (0.3542) |
| **CC(work)** | 0.959 (0.673) |
| **CC(free)** | 0.943 (0.536) |
| **Number of non-hydrogen atoms** | 4539 |
| **macromolecules** | 3927 |
| **ligands / solvent** | 25 / 587 |
| **Protein residues** | 512 |
| **RMS(bonds) / RMS(angles)** | 0.006 / 0.91 |
| **Ramachandran favored (%)** | 98.21 |
| **Ramachandran allowed (%)** | 1.79 |
| **Ramachandran outliers (%)** | 0.00 |
| **Rotamer outliers (%)** | 0.00 |
| **Clashscore** | 2.84 |
| **Average B-factor** | 25.90 |
| **macromolecules** | 23.89 |
| **ligands** | 102.40 |
| **solvent** | 36.09 |
| **Number of TLS groups** | 30 |

**Table S5. Phenix.Table_one for co-crystal of anti-UTag intrabody and UTag (PDB: 8V8F).** Statistics for the highest-resolution shell are shown in parentheses.

| | |
|---|---|
| **Wavelength** | 1.0 |
| **Resolution range** | 45.71 - 2.27 (2.351 - 2.27) |
| **Space group** | P 1 |
| **Unit cell** | 54.467 61.824 85.5767 68.8341 77.8985 71.375 |
| **Total reflections** | 78817 (7454) |
| **Unique reflections** | 42832 (3973) |
| **Multiplicity** | 1.8 (1.9) |
| **Completeness (%)** | 94.23 (87.85) |
| **Mean I/sigma(I)** | 4.77 (0.78) |
| **Wilson B-factor** | 33.85 |
| **R-merge** | 0.2431 (1.059) |
| **R-meas** | 0.3438 (1.498) |
| **R-pim** | 0.2431 (1.059) |
| **CC1/2** | 0.0186 (0.304) |
| **CC*** | 0.191 (0.682) |
| **Reflections used in refinement** | 42688 (3968) |
| **Reflections used for R-free** | 1998 (186) |
| **R-work** | 0.2284 (0.3273) |
| **R-free** | 0.2755 (0.3759) |
| **CC(work)** | 0.944 (0.627) |
| **CC(free)** | 0.906 (0.641) |
| **Number of non-hydrogen atoms** | 7868 |
| **macromolecules** | 7647 |
| **ligands / solvent** | 45 / 176 |
| **Protein residues** | 998 |
| **RMS(bonds) / RMS(angles)** | 0.002 / 0.58 |
| **Ramachandran favored (%)** | 96.71 |
| **Ramachandran allowed (%)** | 3.29 |
| **Ramachandran outliers (%)** | 0.00 |
| **Rotamer outliers (%)** | 0.62 |
| **Clashscore** | 3.66 |
| **Average B-factor** | 40.56 |
| **macromolecules** | 40.56 |
| **ligands** | 57.60 |
| **solvent** | 36.19 |
| **Number of TLS groups** | 27 |

**Table S6. Binding affinity characterization of each tagging system using FRAP.**

| Tagging system | Recovered intensity (Mean ± SEM) | Half-recovery time ($t_{1/2}$, sec) |
|---|---|---|
| Anti-UTag-IB | 84% ± 1% | 177 ± 7 |
| Anti-UTag-IB(ΔCys) | 78% ± 1% | 194 ± 6 |
| Anti-HA-IB | 90% ± 1% | 39 ± 4 |
| Anti-SunTag-IB | 55% ± 2% | N.A. |
| Anti-ALFA-IB | 39% ± 2% | N.A. |

**Table S7. Translation kinetics determined with different tagging systems.**

| Tagging system | $k_i$ ($s^{-1}$) | $k_e$ (aa/s) | Protein length (aa) | Rib. density (%) | No. ribosomes | Rib. distance (aa) |
|---|---|---|---|---|---|---|
| **Anti-UTag-IB** | 0.031 [0.017, 0.047] | 3.11 [2.10, 4.63] | 1968 | 9.0 | 19.8 | 99.6 |
| **Anti-UTag-IB(ΔCys)** | 0.063 [0.052, 0.076] | 5.29 [4.13, 6.56] | 1968 | 10.7 | 23.5 | 83.8 |
| **Anti-SunTag-IB** | 0.034 [0.022, 0.045] | 4.33 [3.01, 5.94] | 2133 | 7.1 | 16.7 | 127.6 |
| **Anti-ALFA-IB** | 0.043 [0.032, 0.053] | 4.84 [3.82, 5.85] | 2016 | 8.0 | 17.8 | 113.0 |

$k_i$ and $k_e$: Parameters calculated with top 5% ranked solutions (shown in brackets).
Protein length: Total length of the reporter protein plus the 24xtag array.
Ribosome density: Percentage of codons occupied by ribosomes at steady state.
Number of ribosomes: Average number of ribosomes simultaneously translating each transcript.
Ribosome distance: Average distance between consecutive ribosomes along the mRNA.

**Table S9. Statistical summary and bootstrap analysis of Rosetta Energy Units (REU) for cysteine-free intrabody variants.**

| Name | Mutations | REU |
|---|---|---|
| G1_mutant_000 | L6L-L81L-C24A-C98A | -770.9 ± 0.7 |
| G1_mutant_002 | L6L-L81L-C24V-C98A | -768.4 ± 1.0 |
| **G1_mutant_009** | **L6I-L81L-C24A-C98A** | **-769.3 ± 0.8** |
| G1_mutant_011 | L6I-L81L-C24V-C98A | -764.5 ± 0.7 |
| G1_mutant_012 | L6I-L81I-C24A-C98A | -768.8 ± 0.9 |
| G2_mutant_004 | L142L-M175I-C161A-C230V | -766.4 ± 0.6 |
| G2_mutant_005 | L142L-M175I-C161V-C230A | -767.8 ± 0.9 |
| G2_mutant_006 | L142L-M175M-C161A-C230A | -763.0 ± 0.5 |
| **G2_mutant_008** | **L142L-M175M-C161V-C230A** | **-770.1 ± 0.7** |
| G2_mutant_017 | L142I-M175M-C161V-C230A | -766.9 ± 0.7 |

Energetic stability was evaluated for the top five variants within each mutation group. The mean and standard deviation (± SD) were calculated from a bootstrap analysis (1,000 iterations), where 20 samples out of 60 total were drawn with replacement in each iteration. Final designs selected for experimental testing based on REU scores and structural intuition are indicated in bold.
